# Translational regulation by RACK1 in astrocytes represses KIR4.1 expression and regulates neuronal activity

**DOI:** 10.1101/2022.07.16.500292

**Authors:** Marc Oudart, Katia Avila-Gutierrez, Clara Moch, Elena Dossi, Giampaolo Milior, Anne-Cécile Boulay, Mathis Gaudey, Julien Moulard, Bérangère Lombard, Damarys Loew, Alexis-Pierre Bemelmans, Nathalie Rouach, Clément Chapat, Martine Cohen-Salmon

## Abstract

The regulation of translation in astrocytes, the main glial cells in the brain, remains poorly characterized. We developed a high-throughput proteomic screen for polysome-associated proteins in astrocytes and focused on the ribosomal protein receptor of activated protein C kinase 1 (RACK1), a critical factor in translational regulation. In astrocyte somata and perisynaptic astrocytic processes (PAPs), RACK1 preferentially bound to a number of mRNAs, including *Kcnj10*, encoding the inward rectifying potassium (K^+^) channel KIR4.1, a critical astrocytic regulator of neurotransmission. By developing an astrocyte-specific, conditional RACK1 knock-out mouse model, we showed that RACK1 repressed the production of KIR4.1 in hippocampal astrocytes and PAPs. Reporter-based assays revealed that RACK1 controlled *Kcnj10* translation through the transcript’s 5’ untranslated region. Upregulation of KIR4.1 in the absence of RACK1 modified the astrocyte territory volume and neuronal activity attenuatin burst frequency and duration in the hippocampus. Hence, astrocytic RACK1 represses KIR4.1 translation and influences neuronal activity.

## Introduction

Astrocytes, the main glial cells in the brain, are large and highly ramified. They project long processes to neurons (perisynaptic astrocytic processes, PAPs) and brain blood vessels (perivascular astrocytic processes, PvAPs) and dynamically regulate synaptic and vascular functions through the expression of specific, polarized molecular repertoires (Cohen-Salmon et al., 2021; Dallerac et al., 2018). There are few data on the mechanisms that regulate translation in astrocytes. Translation is known to be mediated by cis-acting elements, including RNA motifs and secondary structures that influence the binding of trans-acting proteins (also known as RNA-binding proteins, RBPs) (Harvey et al., 2018). A few RBPs have been identified and studied in astrocytes (Blanco-Urrejola et al., 2021; Mazare et al., 2021). For example, *fragile-X* mental retardation protein (FMRP) has been shown to bind and transport mRNAs encoding autism-related signaling proteins and cytoskeletal regulators in radial glial cells (Pilaz et al., 2016). The selective loss of FMRP in astrocytes was shown to dysregulate protein synthesis in general and expression of the glutamate transporter GLT1 in particular (Higashimori et al., 2016). In the mouse, the expression of a pathological form of FMRP (linked to late-onset fragile X syndrome/ataxia syndrome) in astrocytes was found to impair motor performance (Wenzel et al., 2019). More recently, mRNAs enriched in PAPs were shown to contain a larger number of Quaking-binding motifs (Sakers et al., 2021), and inactivation of the cytoplasmic Quaking isoform QKI-6 in astrocytes altered the binding of a subset of mRNAs to ribosomes (Sakers et al., 2021). *Quaking* was also shown to regulate the differentiation of neural stem cells into glial precursor cells by upregulating several genes involved in gliogenesis (Takeuchi et al., 2020). Another general parameter of importance in the regulation of translation is the composition of the translation machinery itself, including ribosomal RNAs (rRNA) and proteins (Gay et al., 2022; Mauro and Matsuda, 2016). This aspect had not previously been studied in astrocytes. Lastly, RNA distribution and local translation are important, highly conserved mechanisms for translational regulation in most morphologically complex cells (Besse and Ephrussi, 2008). We and others have demonstrated that local translation occurs in astrocyte PvAPs and PAPs; this translation might sustain the cells’ molecular and functional polarity (Boulay et al., 2017; Mazare et al., 2020b; Sakers et al., 2017).

To advance our understanding of translation mechanisms in astrocytes, we identified a pool of polysome-associated proteins in astrocytes by combining translating ribosome affinity purification (TRAP) (Mazare et al., 2020a) with mass spectrometry (TRAP-MS). We then focused on receptor of activated protein C kinase 1 (RACK1), a highly conserved eukaryotic protein that is involved in several aspects of translation. RACK1 is positioned at the head of the 40S subunit in the vicinity of the mRNA exit channel (Gallo and Manfrini, 2015; Nilsson et al., 2004). It regulates not only ribosome activities (such as frameshifting and quality-control responses) but also polysome localization and mRNA stability (Ikeuchi and Inada, 2016; Juszkiewicz et al., 2020). In the brain, RACK1 has been mainly described in neurons (Kershner and Welshhans, 2017b) and is involved in local translation and axonal guidance and growth (Kershner and Welshhans, 2017a). The changes in RACK1 expression observed in several neuropathological contexts (such as bipolar disorder (Wang and Friedman, 2001), Alzheimer’s disease (Battaini and Pascale, 2005; Battaini et al., 1999), epilepsy (do Canto et al., 2020; Xu et al., 2015), addiction (McGough et al., 2004), amyotrophic lateral sclerosis (Russo et al., 2017), and Huntington’s disease (Culver et al., 2012)) indicate the importance of the protein’s physiological role in the brain.

In the present study, we demonstrate that RACK1 associates with specific mRNAs, represses the translation of *Kcnj10* mRNA (encoding the inward rectifying K^+^ channel KIR4.1), and regulates neuronal activity.

## Results

### Identification of polysome-associated proteins in astrocytes

We used TRAP-MS to identify polysome-associated proteins in astrocytes **(Fig. 1A)**. Enhanced green fluorescent protein (eGFP)-tagged polysomal complexes were immunopurified from whole brain cytosolic extracts prepared from 2-month-old Aldh1l1:L10a-eGFP transgenic BacTRAP (BT) mice. These animals express the eGFP-tagged ribosomal protein RPL10a specifically in astrocytes (Heiman et al., 2008) **(Fig 1A)**. The same experiment was performed on brain samples from C57/BL6 (wild type) mice, as a control **(Fig 1A)**. A Western blot analysis of immunoprecipitated proteins showed that RPL10a-GFP and the ribosomal protein S6 (RPS6, a component of the 40S ribosomal subunit) were present in BT immunoprecipitates only – demonstrating the efficiency and specificity of TRAP-MS **(Fig. 1B)**. Extracted proteins were characterized using proteomics and quantitative label-free tandem MS (LC-MS-MS) **(Fig 1A, C)**.Three proteins were found only in WT extracts and 139 were found only in BT extracts (fold changes (FC): – or +οο), 61 proteins were enriched in WT extracts (p-value < 0.05; Log2 FC < −1), 106 proteins were detected in both WT and BT extracts (p-value < 0.05; −1 < Log2 FC < 1), and 110 proteins were enriched in BT extracts (p-value < 0.05; Log2 FC > 1) **(Fig. 1C; Table 1; Table S1).** A Gene Ontology (GO) analysis of the 249 proteins enriched or specifically identified in BT immunoprecipitates indicated that most were ribosomal proteins (26%) or RBPs (39.1%) involved in ribosome biogenesis (22.7%) and gene expression (30.9%) (**Fig. 1D)**. We were able to identify a set of polysome-associated proteins in astrocytes.

**Figure 1:**
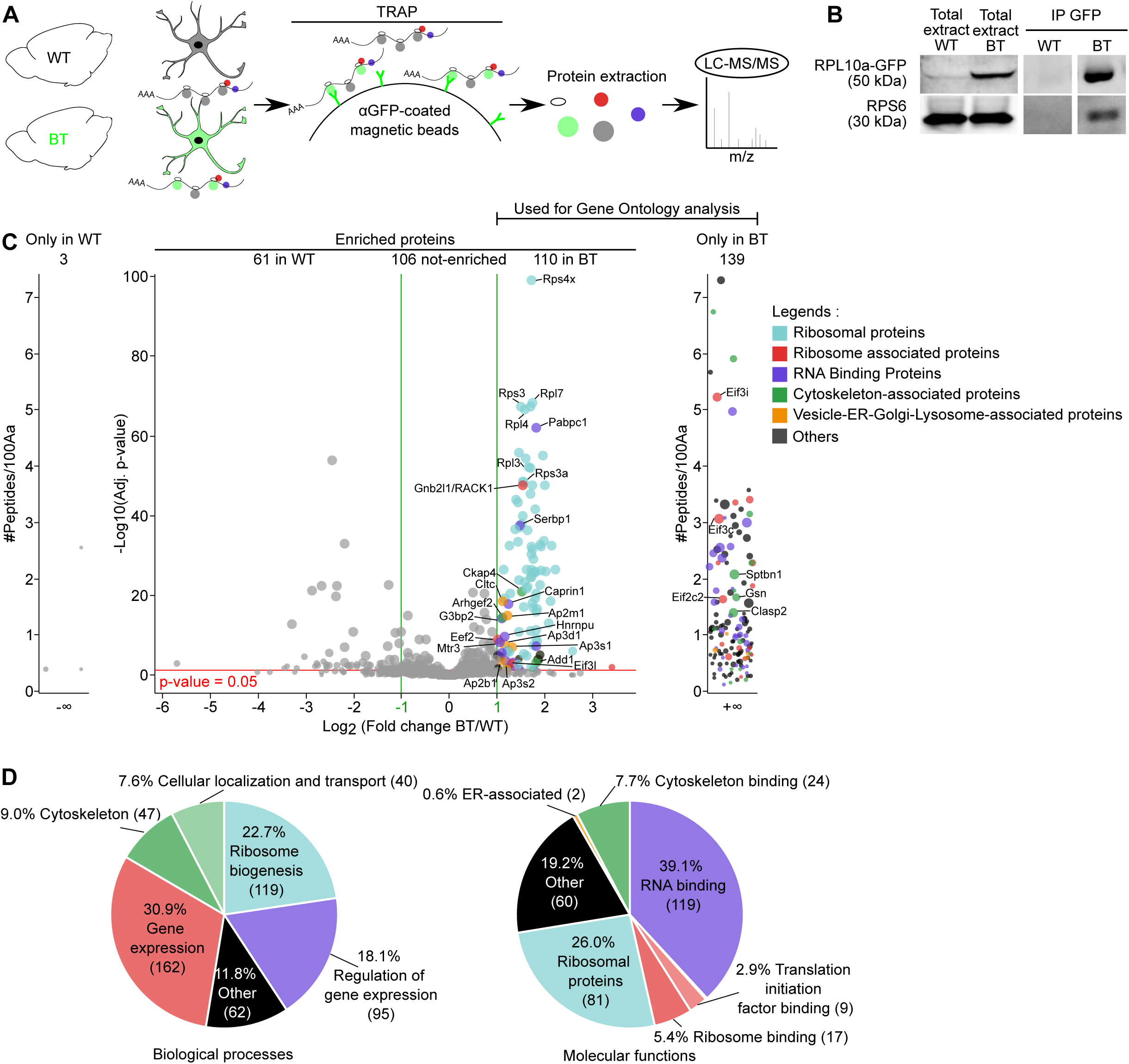
Identification of polysome-associated proteins in astrocytes. **A**. Flowchart of the TRAP-MS analysis on whole brain extracts. Proteins extracted from whole brain in C57/Bl6 WT mice or Aldh1l1:L10a-eGFP (BT) mice were immunoprecipitated by TRAP and analyzed by LC-MS/MS. **B.** Western blot detection of Rpl10a-eGFP and RPS6 in whole brain extracts or TRAP immunoprecipitated proteins (IP-GFP). WT extracts were used as negative controls. **C.** Volcano plot of the TRAP-MS results. Each protein is represented by a dot. The dot size is proportional to the number of peptides identified by LC-MS/MS. Dots for proteins specific to or enriched in BT mice are represented with a color code: ribosomal proteins are given in light blue, with ribosome-associated proteins in red, RNA-binding proteins in purple, cytoskeleton-associated proteins in green, vesicles-ER-Golgi-lysosome-associated proteins in orange, and other proteins in black. Five independent replicates were analyzed (one brain per sample). The protein distribution is represented as the Log2 FC of the BT/WT (x-axis) versus −Log10 adjusted p-value (y axis): Proteins identified only in WT extracts (3 proteins) or only in BT extracts (139 proteins) (FC: – or +∞); Proteins enriched in WT or BT extracts. The threshold for the enrichment in WT or BT extracts is p-value < 0.05 (red line) and Log2 FC > 1 or < −1 (green lines). 61 proteins were enriched in WT extracts (p-value < 0.05; Log2 FC < −1), 106 proteins were detected with a similar abundance in WT and BT extracts (p-value < 0.05, −1 < Log2 FC < 1) and 110 proteins were enriched in BT extracts (p-value < 0.05; Log2 FC > 1). **D.** A GO analysis of the 249 proteins enriched or detected solely in BT extracts (p-value < 0.05; Log2 FC > 1) for biological processes (left) and molecular functions (right). The raw data are given in **Table S1**.

**Table 1:** The most abundant polysome-associated proteins in astrocytes, identified using TRAP-MS. The raw TRAP-MS data for a selection of the most specific (+∞) or enriched (p-value < 0.05; Log2 FC > 1) abundant immunoprecipitated proteins in the BT condition. FC, fold-change. The first six proteins in each GO “molecular function” term are listed.

| Uniprot Accession number | Gene name | Protein description | Log<sub>2</sub> FC (BT/WT) | p-value | Peptides used | MW (kDa) |
| --- | --- | --- | --- | --- | --- | --- |
| **Ribosomal proteins (75)** |  |  |  |  |  |  |
| P62702 | Rps4x | 40S ribosomal protein S4, X isoform | 1.72 | 6.99E-100 | 305 | 29.6 |
| Q9D8E6 | Rpl4 | 60S ribosomal protein L4 | 1.57 | 2.66E-67 | 286 | 47.2 |
| P27659 | Rpl3 | 60S ribosomal protein L3 | 1.65 | 5.67E-53 | 253 | 46.1 |
| P62908 | Rps3 | 40S ribosomal protein S3 | 1.51 | 3.98E-68 | 243 | 26.7 |
| P97351 | Rps3a | 40S ribosomal protein S3a | 1.55 | 3.20E-49 | 240 | 29.9 |
| P14148 | Rpl7 | 60S ribosomal protein L7 | 1.74 | 5.02E-69 | 224 | 31.4 |
| **Ribosomal associated proteins (19)** |  |  |  |  |  |  |
| P68040 | Gnb2l1 | Receptor of activated protein C kinase 1 | 1.54 | 2.43E-48 | 205 | 35.1 |
| P58252 | Eef2 | Elongation factor 2 | 1.02 | 1.24E-09 | 91 | 95.3 |
| Q8R1B4 | Eif3c | Eukaryotic translation initiation factor 3 subunit C | +∞ | NA | 28 | 105.5 |
| Q8QZY1 | Eif3l | Eukaryotic translation initiation factor 3 subunit L | 1.31 | 1.24E-03 | 19 | 66.6 |
| Q9QZD9 | Eif3i | Eukaryotic translation initiation factor 3 subunit I | +∞ | NA | 17 | 36.5 |
| Q8CJG0 | Eif2c2 | Protein argonaute-2 | +∞ | NA | 14 | 97.3 |
| **RNA binding proteins (43)** |  |  |  |  |  |  |
| P29341 | Pabpc1 | Polyadenylate-binding protein 1 | 1.83 | 1.08E-62 | 226 | 70.7 |
| Q9CY58 | Serbp1 | Plasminogen activator inhibitor 1 RNA-binding protein | 1.49 | 2.36E-38 | 127 | 44.7 |
| Q60865 | Caprin1 | Caprin-1 | 1.24 | 1.13E-18 | 78 | 78.2 |
| P97379 | G3bp2 | Ras GTPase-activating protein-binding protein 2 | 1.10 | 5.44E-15 | 76 | 54.1 |
| Q8K310 | Matr3 | Matrin-3 | 1.05 | 5.50E-09 | 47 | 94.6 |
| Q8VEK3 | Hnrnpu | Heterogeneous nuclear ribonucleoprotein U | 1.17 | 2.04E-10 | 46 | 87.9 |
| **Cytoskeleton associated proteins (20)** |  |  |  |  |  |  |
| Q60875 | Arhgef2 | Rho guanine nucleotide exchange factor 2 | 1.12 | 2.96E-15 | 88 | 112.0 |
| Q8BMK4 | Ckap4 | Cytoskeleton-associated protein 4 | 1.52 | 8.75E-22 | 85 | 63.7 |
| Q62261 | Sptbn1 | Spectrin beta chain, non-erythrocytic 1 | +∞ | NA | 49 | 274.2 |
| Q9QYC0 | Add1 | Alpha-adducin | 1.82 | 7.42E-04 | 25 | 80.6 |
| Q8BRT1 | Clasp2 | CLIP-associating protein 2 | +∞ | NA | 18 | 140.7 |
| P13020 | Gsn | Gelsolin | $+\infty$ | NA | 13 | 85.9 |
| **Vesicles - Golgi - Lysosomes associated proteins (10)** |  |  |  |  |  |  |
| Q68FD5 | Cltc | Clathrin heavy chain 1 | 1.11 | 2.28E-19 | 269 | 191.6 |
| O54774 | Ap3d1 | AP-3 complex subunit delta-1 | 1.16 | 2.17E-08 | 125 | 135.1 |
| P84091 | Ap2m1 | AP-2 complex subunit mu | 1.23 | 1.36E-15 | 79 | 49.7 |
| Q9DBG3 | Ap2b1 | AP-2 complex subunit beta | 1.09 | 1.60E-04 | 39 | 104.6 |
| Q9DCR2 | Ap3s1 | AP-3 complex subunit sigma-1 | 1.31 | 7.88E-08 | 32 | 21.7 |
| Q8BSZ2 | Ap3s2 | AP-3 complex subunit sigma-2 | 1.19 | 3.34E-02 | 19 | 22.0 |

### RACK1 associates with polysomes in astrocytes

Among the ribosome-associated proteins preferentially extracted with TRAP-MS, we focused on RACK1. This protein binds to the small ribosomal subunit 40S and has a key role in the translation of capped, polyadenylated mRNAs (Johnson et al., 2019) (**Fig. 2A**). RACK1’s role in astrocytes had not been assessed previously. However, we recently identified *Gnb2l1* mRNA (encoding RACK1) as one of the most highly enriched, translated mRNAs in PAPs; this finding suggested that RACK1 has an important role at this cellular interface (Mazare et al., 2020b); A Western blot analysis of TRAP-MS immunoprecipitated proteins showed that RACK1 was specifically detected in the BT condition and thus confirmed the co-immunoprecipitation of RACK1 with astrocytic polysomes (**Fig. 2B**). We next characterized RACK1 expression in astrocytes, with a focus on samples from hippocampus. We used fluorescent *in situ* hybridization (FISH) to detect *Gnb2l1* mRNAs in astrocytes **(Fig. 2C)**. Since *Gnb2l1* is ubiquitous, we immunolabeled astrocytes for GFAP. The *Gnb2l1* FISH dots localized on GFAP processes were identified using our recently developed *Astrodot* protocol (Oudart et al., 2020). In line with our previous results, *Gnb2l1* mRNAs were detected somewhat in astrocyte somata but mainly in astrocytic processes (Mazare et al., 2020b) **(Fig. 2C)**. We next performed immunofluorescence imaging of RACK1 on hippocampal sections **(Fig. 2D)**. RACK1 was clearly detected in neurons and GFAP-immunolabeled astrocytic soma and processes (**Fig. 2D**). These results suggested that RACK1 is expressed in astrocytes and associates with astrocytic polysomes.

**Figure 2:**
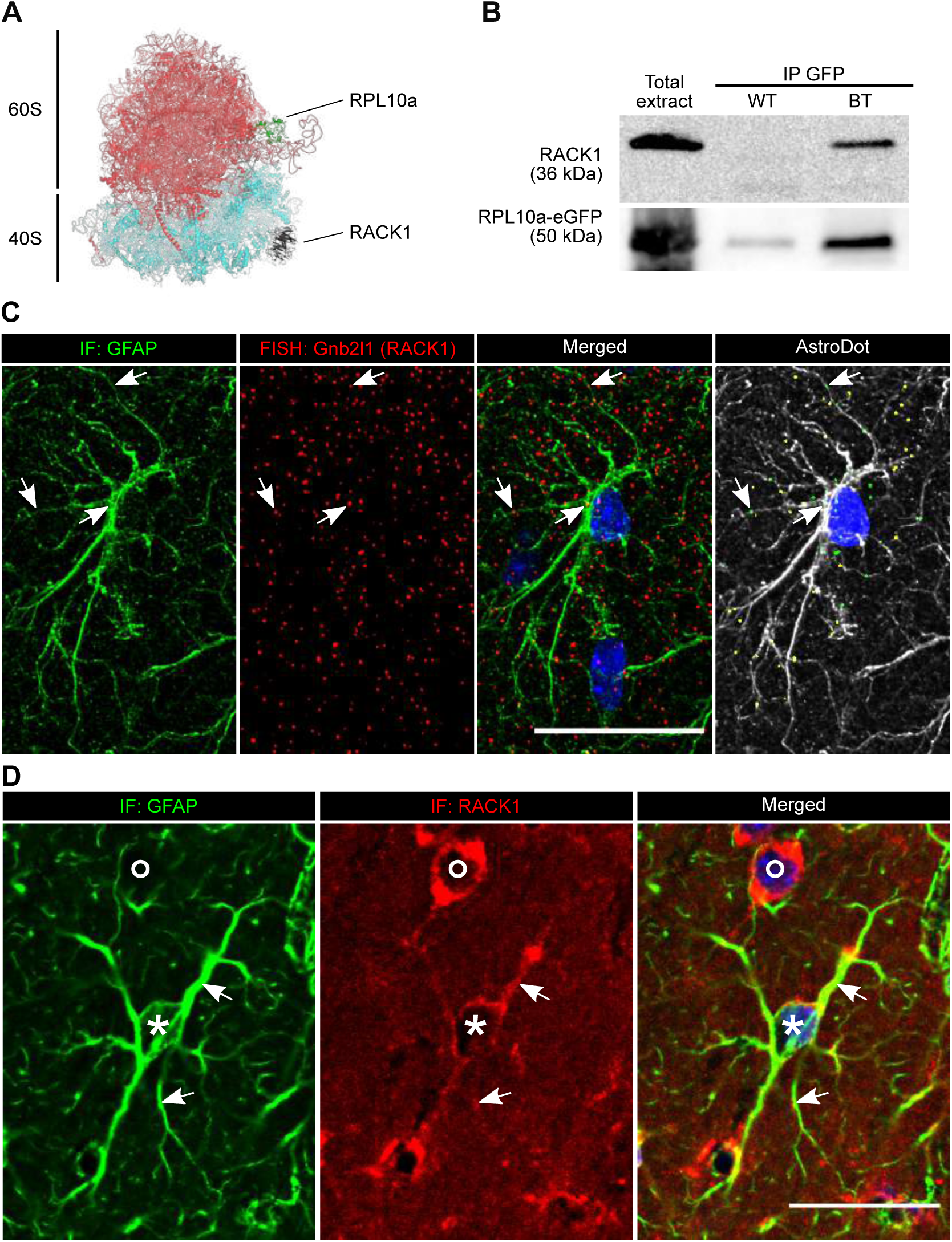
RACK1 is associated with ribosomes in astrocytes. **A.** Representation of the human 80S ribosome, generated with PyMol software (https://pymol.org/2/, PyMol version 2.3.4, Python 3.7) on the basis of the high-resolution cryo-EM structure (Natchiar et al., 2017). RPL10a is shown in green, and RACK1 is shown in black. **B.** Western blot detection of RACK1 and RPl10a-GFP in whole brain protein extracts and in TRAP-MS extracts (IP GFP) in BT conditions. WT extracts were used as negative controls. **C.** FISH detection of *Gnb2l1* mRNAs encoding RACK1 in hippocampal astrocytes immunolabeled for GFAP. **From left to right:** Confocal microscopy image of a GFAP-immunolabeled astrocyte (in green); FISH detection of *Gnb2l1* mRNAs (red dots); merged image; *AstroDot* analysis of *Gnb2l1* mRNA located on GFAP-positive processes. The green dots are located in the soma or in GFAP-immunolabeled large processes; the yellow dots are located in GFAP-immunolabeled fine processes. **D.** Confocal images of RACK1 immunofluorescence detection (in red) in the hippocampus. Astrocytes (*) are co-immunolabeled for GFAP (in green). A neuron (°) is also labelled for RACK1. Scale bar: 20 µm.

### RACK1 associates with specific mRNAs in astrocytes and in PAPs

We next determined which mRNAs were associated with RACK1 in astrocytes via mRNA immunoprecipitation (using a RACK1-specific antibody) of whole brain astrocytic cytoplasmic extracts prepared from 2-month-old mice. We first checked the efficiency of RACK1 immunoprecipitation on Western blots. Increasing levels of anti-RACK1 antibody indeed immunoprecipitated higher levels of RACK1 and the small subunit ribosomal protein RPS6 (**Fig. 3A**). We then repeated the experiment with the optimal quantity of RACK1 antibody, extracted the immunoprecipitated mRNAs, and analyzed them with qPCRs (**Fig. 3B**). Nonspecific mouse immunoglobulins G (IgG) were used as a negative control (**Fig. 3B**). Extracts prepared from whole brain were immunoprecipitated (**Fig. 3C**). RACK1 was present in astrocyte processes (**Fig. 2D**) and was preferentially translated in hippocampal PAPs (Mazare et al., 2020b). We therefore also immunoprecipitated RACK1 in synaptogliosome preparations consisting of PAPs attached to synaptic neuronal membranes (Carney et al., 2014) (**Fig. 3C’**). Since RACK1 is ubiquitously expressed in the brain, we limited our analysis on a selection of astrocyte-specific mRNAs and focused on those detected previously in PAPs (Mazare et al., 2020b), such as *Kcnj10*, encoding the inward rectifying K^+^ channel KIR4.1, *Slc1a2*, encoding the glutamate transporter GLT1, *Aqp4*, encoding the water channel aquaporin 4, *Slc1a3*, encoding the glutamate transporter GLAST, and *Gja1* and *Gjb6*, encoding the gap junction proteins connexin 43 and 30, respectively. With the exception of *Gjb6,* all the tested mRNAs were immunoprecipitated more significantly by RACK1 than by IgG in whole brain (**Fig. 3C)** and synaptogliosome extracts (**Fig. 3C’**). These results were probably influenced by the level of polysomal mRNAs in astrocytes and PAPs. We therefore determined the level of each polysomal mRNA in astrocytes and PAPs by performing TRAP and qPCRs on whole-brain extracts from 2-month-old BT mice (**Fig. 3D, E**) or on synaptogliosome extracts (**Fig. 3D, E’**). The mean value for each mRNA was then used to normalize the quantity of RACK1-immunoprecipitated mRNA (**Fig. 3F)** in whole brain (**Fig. 3G**) and in synaptogliosomes (**Fig. 3G’**). The results of these experiments suggested that *Slc1a2* and *Kcnj10* were preferentially associated with RACK1 in astrocytes (**Fig 3G**) and in PAPs (**Fig. 3G’**).

**Figure 3:**
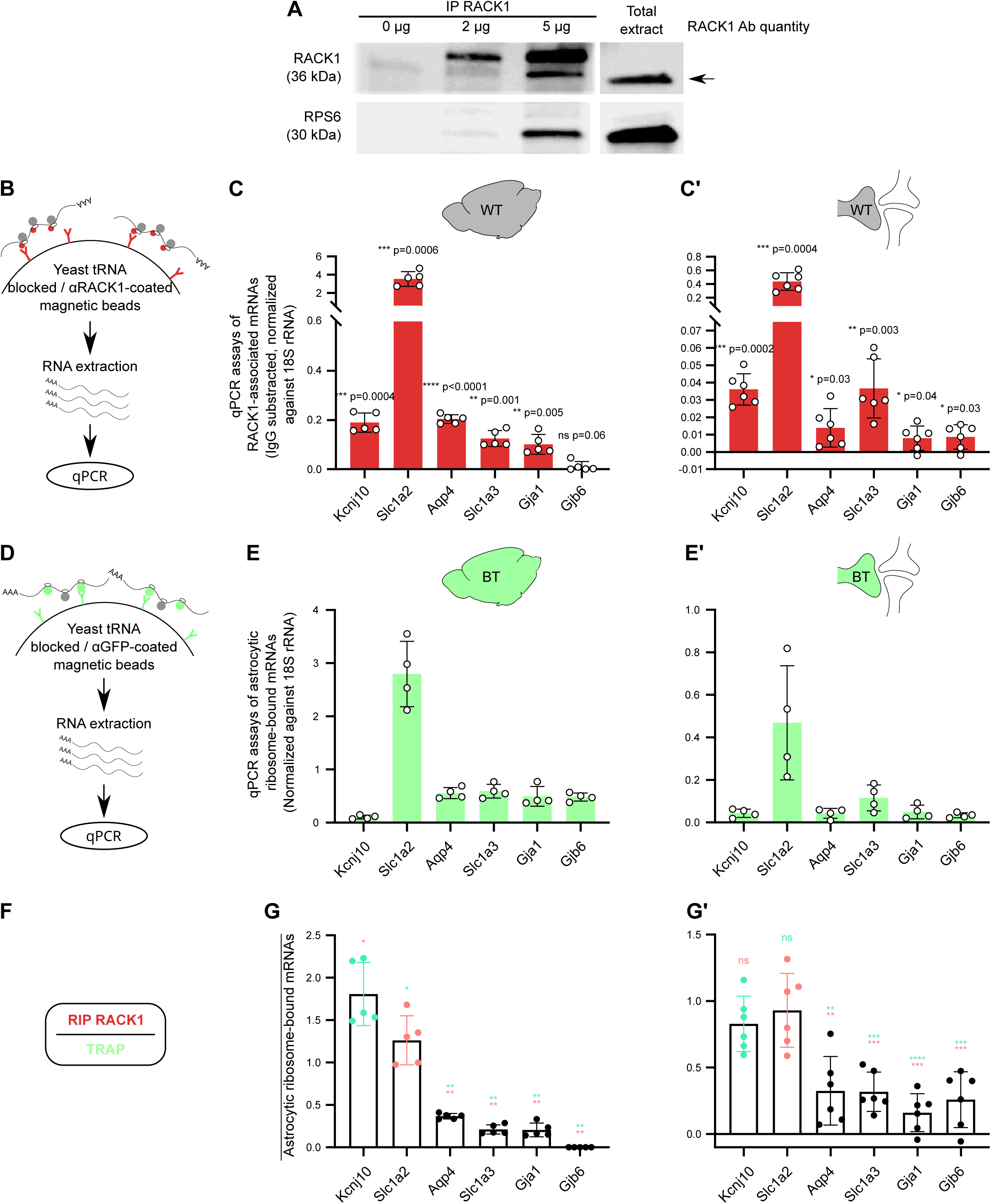
RACK1 associates with specific mRNAs in astrocytes and PAPs. **A.** Western blot analysis of RACK1 immunoprecipitation in whole brain extracts from 2-month-old mice. Increasing quantities of RACK1 antibodies (0, 2, and 5 µg) were used. *Lower panel:* RPS6 is also detected in RACK1-immunoprecipitated proteins. **B.** Flowchart of RNA immunoprecipitation using anti-RACK1 antibodies (in red) on whole brain extracts (**C**) or synaptogliosome extracts (**C’**) prepared from 2-month-old WT mice. Red dots on ribosomes represent RACK1. Immunoprecipitated RNAs were purified and screened (in qPCR assays) for a selection of astrocyte-specific mRNAs. IgG-subtracted signals were normalized against rRNA *18S.* The data are quoted as the mean ± SD (N=5 or 6 samples; 1 mouse brain per sample); one-sample t-test vs. 0 (except for *Gjb6* whole brain experiment, One-sample Wilcoxon test). **D.** Flowchart of polysomal immunoprecipitation (TRAP) using anti GFP antibodies (in green) on whole brain (**E**) or synaptogliosomes extracts (**E’**) prepared from 2-month-old BT mice. Immunoprecipitated RNAs are purified and analyzed by qPCR for a selection of astrocyte specific mRNAs. Signals were normalized against rRNA *18S.* The data are quoted as the mean ± SD (N=4 samples; 1 mouse brain per sample). **F.** Flowchart of the normalization of the RACK1 immunoprecipitation against mean TRAP values. Ratios were calculated for experiments on whole brain (**G**) or synaptogliosomes (**G’**). The data are quoted as the mean ± SD (N=5 or 6 samples; 1 mouse brain per sample). P values are indicated in green when *Kncj10* results was the reference and in red when *Slc1a2* results was the reference. Two-tailed unpaired t-test or a two-tailed Mann-Whitney test. ns, not significant (p>0.05); *, p<0.05; **, p<0.01; ***, p<0.001; ****, p<0.0001. The raw data are presented in **Table S2**.

Taken as a whole, these results suggested that RACK1 associates preferentially with specific mRNAs in astrocytes and in PAPs.

### RACK1 represses the expression of *Kcnj10* in astrocytes

To gain insights into RACK1’s function in astrocytes, we generated a RACK1 conditional knock-out mouse model (RACK1 cKO) by crossing RACK1 fl/fl mice with Aldh1L1-CreERT2 mice (**Fig. 4A**). Two month-old Aldh1L-CreERT2: RACK1 fl/fl mice were injected with tamoxifen, to induce RACK1 KO in astrocytes (**Fig 4A**). *Gnb2l1* KO in astrocytes was confirmed by PCRs on DNA extracted from whole brain (**Fig. 4B**) and by immunofluorescence assays of hippocampal sections (**Fig. 4C**). In the cKO mice, RACK1 was detected in pyramidal layer neurons but not in astrocytes immunolabelled for GFAP (**Fig. 4C**). We next sought to determine the impact of RACK1 KO in astrocytes on the level of GLT1 and KIR4.1 in whole hippocampus or hippocampal synaptogliosome protein extracts from RACK1 fl/fl and RACK1 cKO mice (**Fig. 4D**). Interestingly, the various extracts did not differ significantly with regard to the level of GLT1 but the level of KIR4.1 was significantly higher in the two extracts from RACK1 cKO mice (**Fig. 4D**).

**Figure 4:**
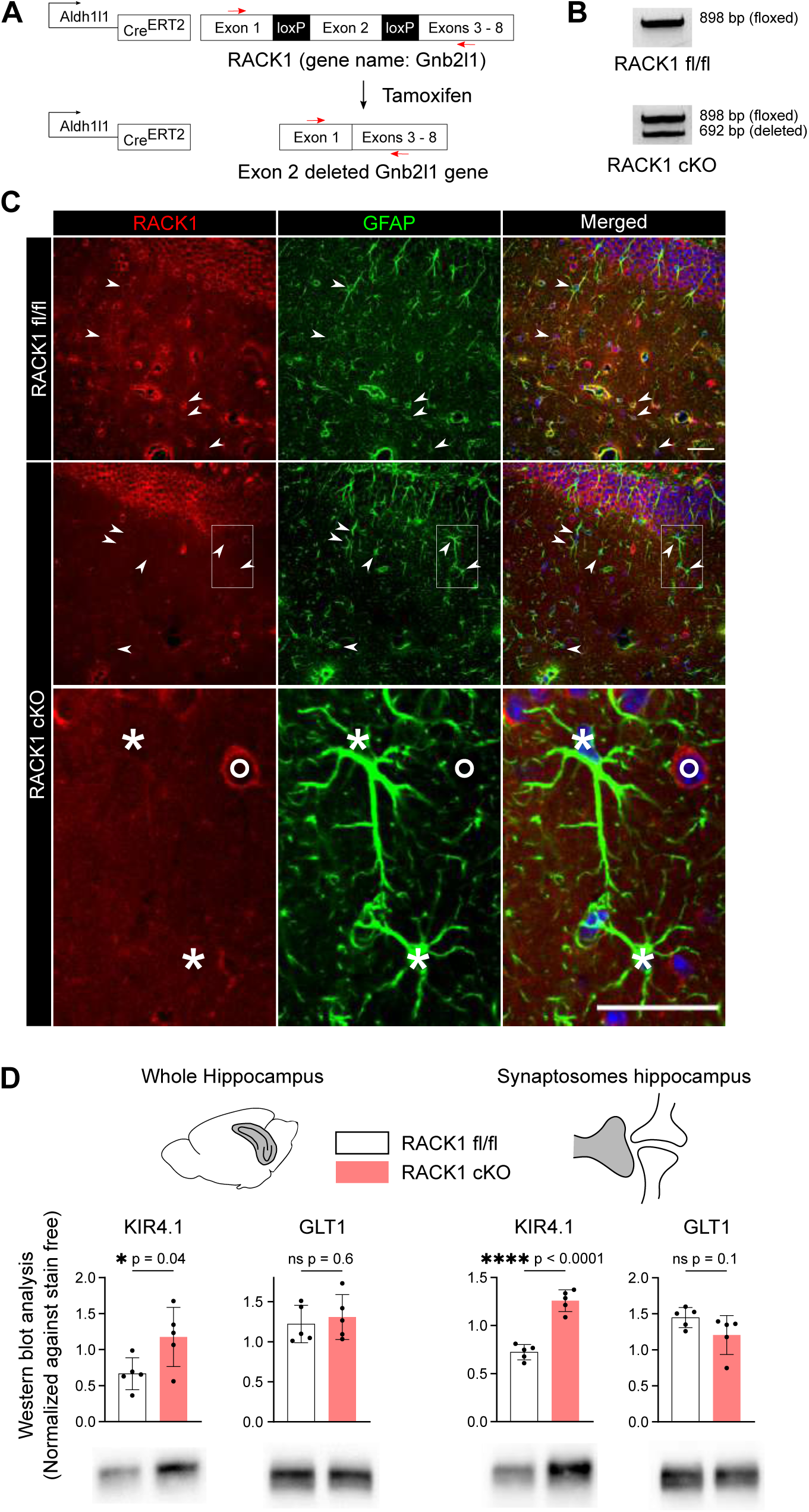
RACK1 KO in astrocytes leads to higher levels of KIR4.1 in astrocyte somata and PAPs. **A.** Generation of a mouse line with RACK1 knocked out in astrocytes. Schematic representation of the RACK1 fl/fl and Aldh1l1-Cre/ERT2 alleles. Deletion of exon 2 in *Gnb2l1* (the gene coding for RACK1) is induced in astrocytes by tamoxifen injection; this results in a frameshift and the premature termination of *Gnb2l1* translation. **B.** PCR assays for *Gnb2l1* KO in brain DNA from RACK1 fl/fl or Aldh1L-CreERT2/ RACK1 fl/fl tamixofen-injected mice (RACK1 cKO). Primers are indicated in (**A**) by red arrows. The 898 base-pair (bp) band corresponds to the floxed allele. The 672 bp band corresponds to the exon2-deleted allele**. C.** Confocal images of RACK1 immunofluorescence detection (in red) in the hippocampus in RACK1 fl/fl and RACK1 cKO mice. Astrocytes are co-immunolabeled for GFAP (in green). The lower panel gives a higher magnification view of the boxed area in the RACK1 cKO images, which shows that RACK1 is specifically depleted in astrocytes (*) and is still expressed by neurons (°). **D.** Western blot detection and analysis of KIR4.1 and GLT-1 in protein extracts from whole hippocampus or synaptogliosomes purified from RACK1 fl/fl or RACK1 cKO mice. The data are quoted as the mean ± SD (N=5 samples per genotype; 1 mouse per sample); two-tailed unpaired ttest. ns, not significant, p-value>0.05; *, p<0.05; **, p<0.01; ***, p<0.001; ****, p<0.0001. The raw data are presented in **Table S2**.

These results demonstrated that RACK1 deficiency in astrocytes led to a higher level of KIR4.1 in whole astrocytes and in PAPs.

### *In cellulo* translational control by RACK1 depends on the 5’ untranslated region (5’UTR) of ***Kcnj10***

RACK1 is an essential factor in translation and in ribosome quality control. It senses ribosome stalling on rare codons (such as CGA, coding for arginine, and AAA, coding for lysine) and can cause translation elongation to pause or abort. The absence of RACK1 results in the more frequent translation of mRNAs with stalling sequences and eventually the accumulation of peptides with frameshifts (Juszkiewicz et al., 2020). Since we observed RACK1-dependent downregulation of *Kcnj10*, we first hypothesized that the *Kcnj10* gene’s coding sequence (CDS) is subject to a stalling event that can only be resolved by RACK1. To test this hypothesis, we generated Human Embryonic Kidney 293T (HEK293T) cells in which RACK1 expression was disrupted through a CRISPR/Cas9-based strategy (RACK1^KO^ cells; **Fig. 5A**). We then designed a dual fluorescence reporter system in which GFP and mCherry fluorescent proteins were expressed in-frame from a single mRNA and were separated by the *Kcnj10* CDS (**Fig. 5B**). The *Kcnj10* CDS was insulated with viral P2A sequences, at which ribosomes skip the formation of a peptide bond without interrupting elongation (Lin et al., 2013). Complete translation of this cassette generates three proteins (GFP, mKIR4.1, and mCherry), and the presence of any stall-inducing sequences in *Kcnj10* CDS would modify the translation rate prior to mCherry synthesis and would thus result in a sub-stoichiometric mCherry:GFP ratio. A reporter without *Kcnj10* CDS served as a negative control. The positive control was a reporter in which *Kcnj10* CDS had been replaced by a sequence containing a stretch of consecutive lysine AAA codons (termed K20) and that was known to induce ribosome stalling (**Fig.S1A**) (Juszkiewicz and Hegde, 2017). These reporters were expressed in WT and RACK1^KO^ cells, and the mCherry:GFP ratio was measured at the single-cell level using fluorescence-activated cell sorting (**Fig. 5C, Fig. S1B**). We found that RACK1^KO^ cells displayed a robust elevation of the mCherry level expressed downstream of the K20 sequence, confirming that loss of RACK1 impairs ribosome stalling (**Fig. S1B, C**). In contrast, the absence of RACK1 did not modify the mCherry/GFP ratio of the reporter cassette containing *Kcnj10* CDS, when compared with WT cells (**Fig. 5C, D**). Taken as a whole, these data indicate that the sensitivity of the *Kcnj10* mRNA with regard to RACK1 is not mediated by a RACK1-modulated ribosomal event involving its CDS.

**Figure 5:**
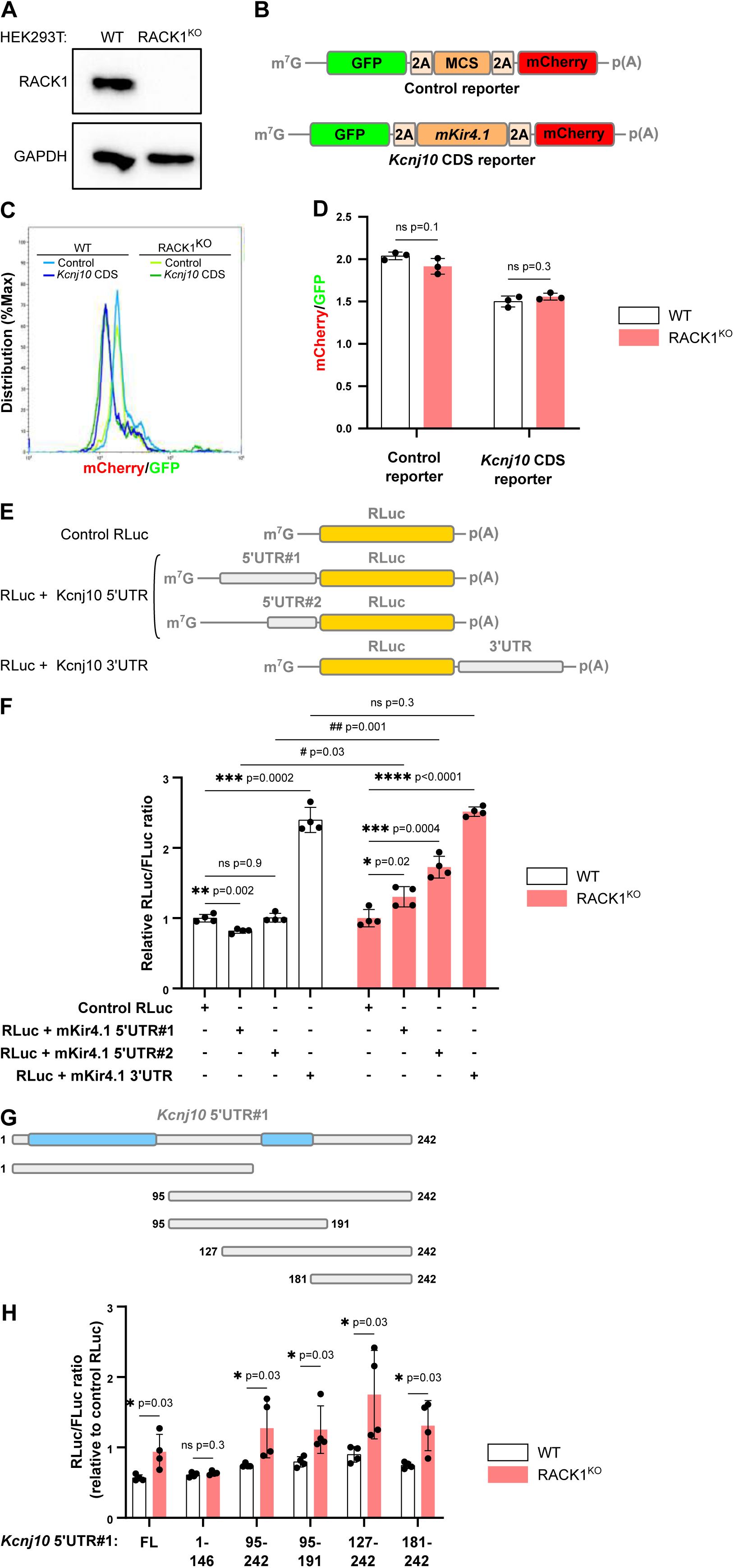
The 5’UTR of *Kcnj10* confers RACK1 sensitivity *in vitro*. **A.** CRISPR/CAS9-based KO of RACK1 in HEK293T cells. The Western blot for the indicated proteins was performed using WT and RACK1^KO^ cell extracts. **B.** Topology of the reporters for flow cytometry analysis of *Kcnj10* mRNA translation. The constructs contain GFP and mCherry separated by a multiple cloning site (into which the *Kcnj10* CDS had been inserted) and two viral 2A sequences (at which ribosomes skip formation of a peptide bond, without interrupting chain elongation). **C.** Representative flow-cytometry-based assay of the fluorescence ratio in WT or RACK1^KO^ HEK293T cells transfected by the constructs described in (**B**). **D.** A histogram of the data, presented as the mean ± SD (N = 3); Two-tailed t-test with Welch’s correction. **E**. Schematic representation of the *Renilla* luciferase (RLuc) reporter constructs harboring the 5’UTR and/or 3’UTR of mouse *Kcnj10* mRNA. These sequences were inserted in the psiCHECK2 vector, which also encodes the Firefly luciferase (Fluc). The “control” RLuc was generated by transfecting the empty psiCHECK2 vector. **F**. WT and RACK1^KO^ HEK293T cells were transfected with the reporters described in (**E**). Luciferase activity was measured 24 h after transfection. RLuc values were normalized against FLuc levels, and a ratio was calculated for each *Kcnj10* reporter relative to the empty psiCHECK2 reporter (value set to 1) for each population. **F.** A histogram of the data, presented as the mean ± SD (N = 4); unpaired Mann-Whitney test or unpaired t-test. **G.** Schematic representation of the truncated versions of the *Kcnj10* 5’UTR inserted in the RLuc reporter. Blue boxes indicated GC-rich regions (see **Fig.** S**2B**). Plasmids were transfected in WT and RACK1^KO^ HEK293T cells. For each experiment, the ratio was calculated for each *Kcnj10* reporter relative to the empty psiCHECK2 reporter (value set to 1). **H**. A histogram of the data, presented as the mean ± SD (N = 4); unpaired Mann-Whitney test. ns, not significant, pvalue>0.05; *, p<0.05; **, p<0.01; ***, p<0.001; ****, p<0.0001. The raw data are presented in **Table** S**2**.

We next sought to determine whether *Kcnj10*’s sensitivity to RACK1 was conferred by its 5’UTR. Two distinct 5’UTRs have been reported for *Kcnj10* in the mouse (NM_001039484.1 and AB039879.1, hereafter referred to respectively as 5’UTR#1 and 5’UTR#2). 5’UTR#1 is composed of a G/C-rich first half (region 1-104, not found in other mammalian *Kcnj10* orthologs) and a highly conserved second half (the 147-242 region which is shared with 5’UTR#2) (**Fig. S2A, Fig. 5E**). 5’UTR#1 and 5’UTR#2 were inserted in the psiCHECK-2 luciferase reporter vector downstream of the *Renilla* luciferase (RLuc) CDS (**Fig. 5E**). A reporter harboring the 3’UTR of *Kcnj10* upstream of RLuc was also constructed as a control. These reporters were transfected into WT and RACK1^KO^ HEK293T cells, and the effect of RACK1 loss on RLuc activity for each UTR construct was calculated relative to an empty psiCHECK2 reporter level (control RLuc). RLuc activity was normalized against the activity of the co-expressed FLuc (**Fig. 5F**). We detected significantly greater RLuc activity when the RLuc reporters harboring *Kcnj10* 5’UTRs were expressed in RACK1^KO^ cells (relative to expression in WT cells) (**Fig. 5F**). In contrast, no difference between WT and RACK1^KO^ cells was observed for the RLuc-Kcnj10 3’UTR construct – indicating that translational control by RACK1 is mediated by *Kcnj10* 5’UTRs (**Fig. 5F**). A qPCR analysis did not show significant differences in RLuc mRNA levels under any conditions, which confirmed that the *Kcnj10* 5’UTR-mediated effect on RLuc activity was post-transcriptional (**Fig. S3A,B**). Since 5’UTR#1 and 5’UTR#2 share a common 96-nucleotide region (**Fig. S2B**), we hypothesized that this sequence confers RACK1-dependent translation control. To test this hypothesis, we truncated the 242-nucleotide-long *Kcnj10* 5’UTR#1 into five overlapping fragments, which were inserted upstream of the RLuc sequence and expressed in WT and RACK1^KO^ cells (**Fig. 5G**). We found that both the 127-242 region and the shorter 181-242 region were sufficient to increase the RLuc activity in RACK1^KO^ cells (relative to WT cells), whereas the first half (region 1-146) did not confer RACK1 sensitivity (**Fig. 5G, H**).

Taken as a whole, these data demonstrate that RACK1’s control over the translation of *Kcnj10* mRNA depends on the *Kcnj10* 5’UTR rather than its CDS or 3’UTR.

### RACK1 regulates astrocyte volume

KIR4.1 is a weakly inwardly rectifying K^+^ channel that confers astrocytes with high K^+^ conductance. K^+^ influx into astrocytes is thought to be coupled to water intake, leading to transient or prolonged swelling (MacVicar et al., 2002; Risher et al., 2009). Thus, elevation of KIR4.1 levels in RACK1 cKO might be associated with a greater hippocampal astrocyte volume. We used an adeno-associated virus (AAV) bearing the gfaABC1D synthetic promoter (derived from Gfap; (Lee et al., 2008)) to drive the expression of the fluorescent protein tdTomato in astrocytes. AAVs were injected into the CA1 region of the dorsal hippocampus of adult mice (**Fig. 6A**). A three-dimensional (3D) analysis was performed on sparse labelled CA1 hippocampal astrocytes from 2-month-old WT and RACK1 cKO mice (**Fig. 6B-F**). RACK1 cKO astrocytes had a larger territory volume (**Fig. 6D**) and longer distal processes **(Fig. 6F)** than RACK1 fl/fl astrocytes.

**Figure 6:**
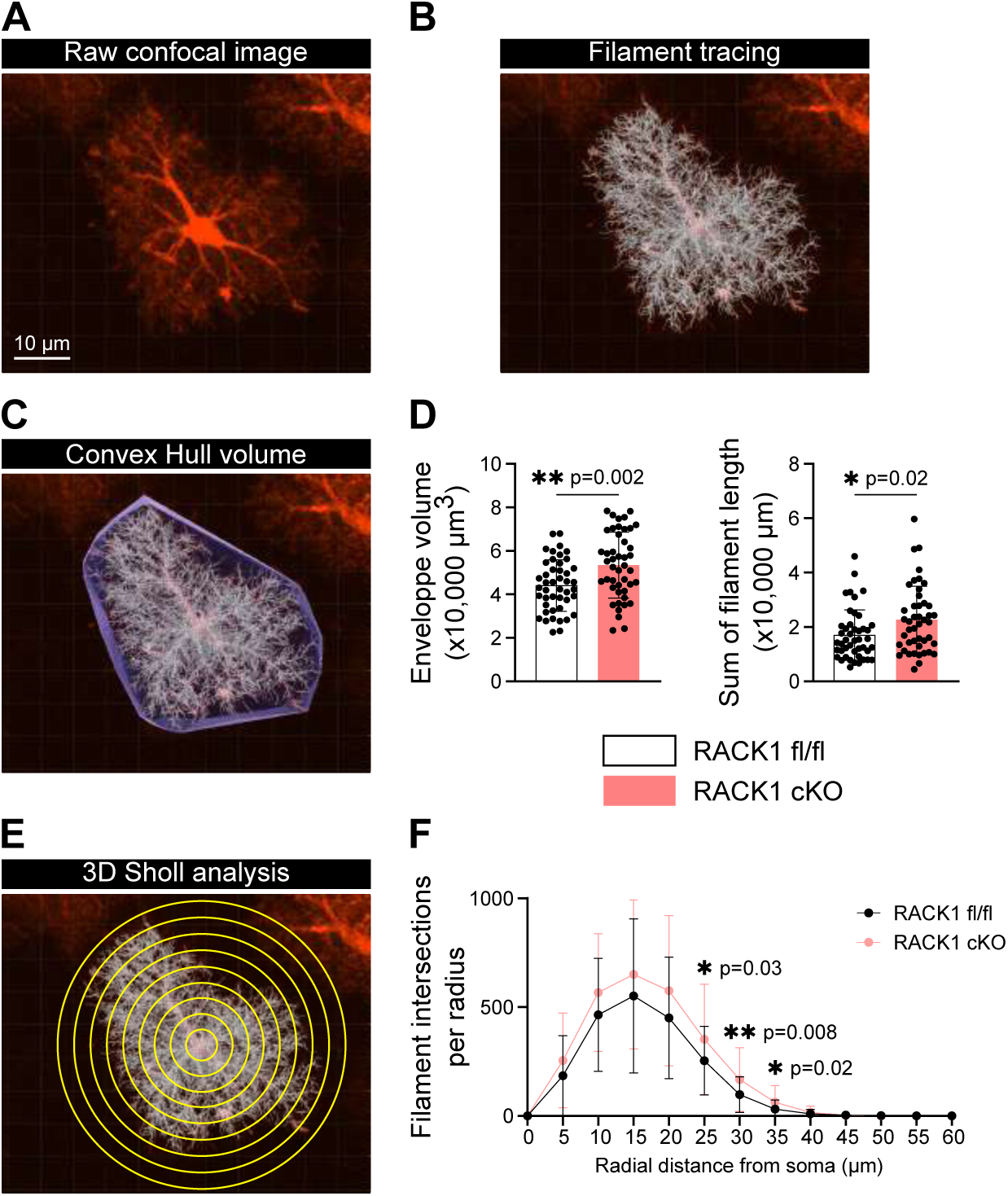
RACK1 regulates astrocyte volume. **A.** Raw confocal image of an isolated CA1 astrocyte expressing tdTomato. **B-F**. Imaris analysis: filament tracing (**B**); convex hull volume (**C**); a 3D Sholl analysis (**E**). **D.** Mean territory volume and filament length of RACK1 fl/fl and RACK1 cKO astrocytes. A histogram of the data, presented as the mean ± SD (N = 4 mice per genotype; 45 astrocytes); two-tailed *t-test*. **F.** A Sholl analysis of the ramification complexity of RACK1 fl/fl and RACK1 cKO astrocytes. Two-way analysis of variance. *, p<0.05; **, p<0.01. The raw data are presented in **Table** S**2**.

These results indicated that astroglial RACK1 is required for a correct hippocampal astrocyte territory volume.

### RACK1 regulates neuronal activity

During their activity, neurons release large amounts of K^+^ at the synapses. The K^+^ is rapidly taken up by astrocytic KIR4.1 and is redistributed across the astrocytic network. This astrocytic K^+^ clearance mechanism maintains perisynaptic homeostasis and prevents neuronal hyperexcitability. Since we had shown that RACK1 cKO astrocytes contained high levels of KIR4.1, we sought to determine whether this change alters basal excitatory synaptic transmission. To this end, we stimulated CA1 Schaffer collateral (SC) synapses in acute hippocampal slices and thus evoked α-amino-3-hydroxy-5-methyl-4-isoxazolepropionic acid receptor (AMPAR)-mediated field excitatory postsynaptic potentials (fEPSPs) (**Fig. 7A**). The size of the presynaptic fiber volley (the input) was compared with the slope of the fEPSP (the output). RACK1 cKO mice and control RACK1 fl/fl mice did not differ with regard to basal synaptic transmission (**Fig. 7B**). We next investigated the effect of the KIR4.1 K^+^ channel blocker VU0134992 (30 µM) and found a similar overall decrease in excitatory synaptic transmission in both RACK1 fl/fl and RACK1 cKO animals after 20 minutes of application (**Fig. 7B**). These results indicate that the elevated expression of astrocytic KIR4.1 K^+^ channels in RACK1 cKO mice does not modify hippocampal basal excitatory synaptic transmission evoked in the hippocampal CA1 region.

**Figure 7:**
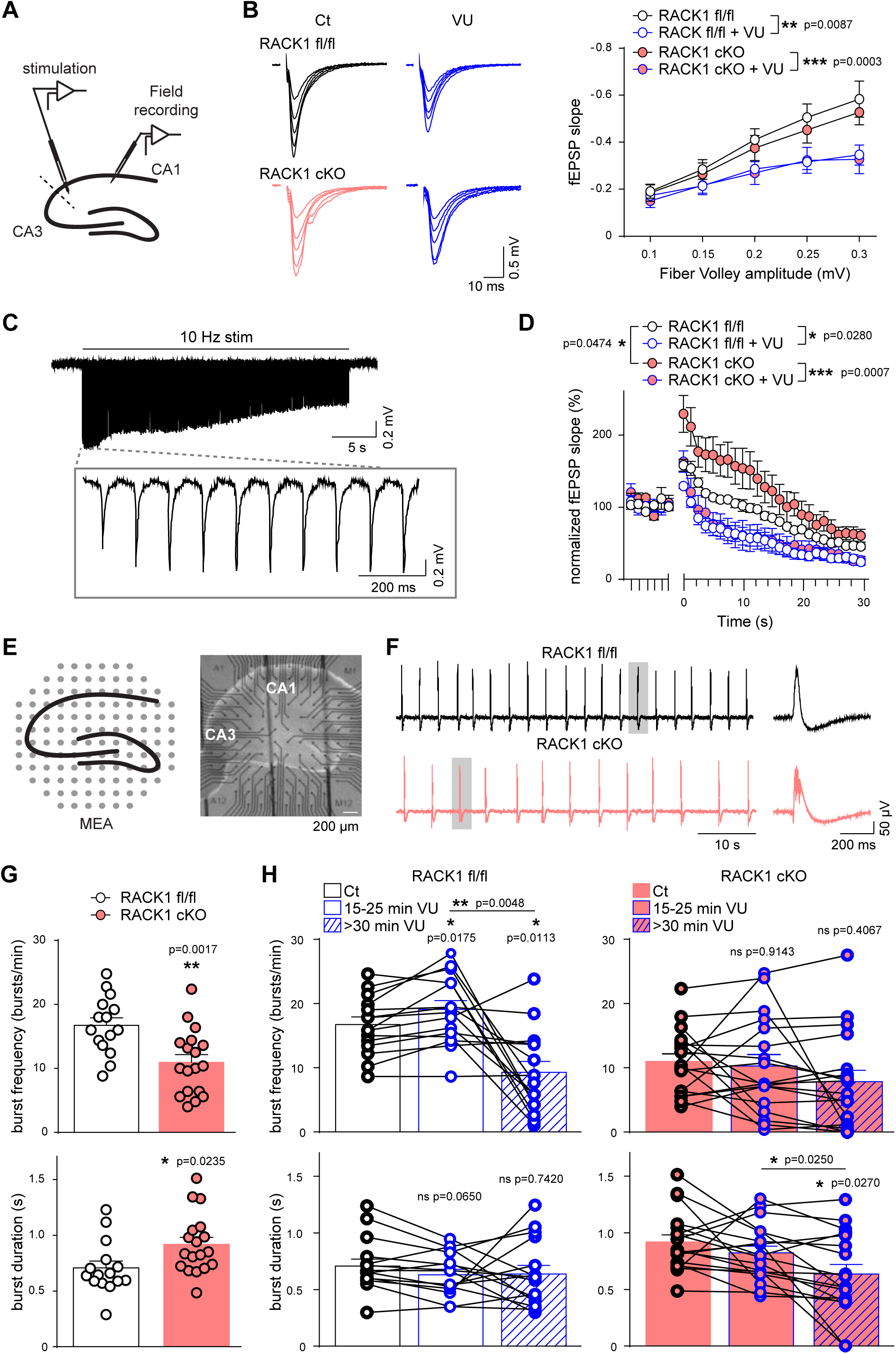
RACK1 KO in astrocytes does not affect basal excitatory synaptic transmission but does alter network population activity and neuronal responses to intense stimulation. **A.** Schematic representation of electrode positions used to record field excitatory postsynaptic potentials (fEPSP) evoked by Schaffer collateral (SC) stimulation in the CA1 region of hippocampal slices. **B.** Input-output curves for basal synaptic transmission. Left, representative recordings in RACK1 fl/fl mice (black) and RACK1 cKO mice before (pink) and after (blue) application of a Kir 4.1 antagonist (VU0134992). Scale bars: 10 ms, 0.5 mV. Right, quantification of the fEPSP slope for different fiber volley amplitudes after SC stimulation. (RACK1 fl/fl: n=5 slices from 4 mice; p=0.0087; RACK1 cKO: n=5 slices from 5 mice; repeated measures two-way ANOVA with Sidak’s correction for multiple comparisons). **C.** Top: a representative recording of fEPSPs evoked by repetitive stimulation (10 Hz, 30 s) of CA1 SCs in RACK1 fl/fl mice under control conditions. Scale bars: 5 s, 0.2 mV. Bottom: enlarged view of fEPSPs evoked by the first 10 stimuli. Scale bars: 200 ms, 0.2 mV. **D.** Quantification of changes in the fEPSP slope induced by 10 Hz stimulation relative to responses measured before the onset of stimulation (baseline responses) in RACK1 fl/fl mice (white filled dots) and in RACK1 cKO mice (pink-filled dots) before (black) and after (blue) application of VU0134992 (RACK1 fl/fl: n=5 from 5 mice; RACK1 cKO: n = 6 slices from 4 mice; repeated measures two-way ANOVA with Sidak’s correction for multiple comparisons). **E.** Schematic representation (left) and picture (right) of a hippocampal slice placed on a multielectrode array (MEA). Scale bar: 200 µm. **F.** Representative MEA recordings of burst activity induced in hippocampal slices of RACK1 fl/fl (black) and RACK1 cKO (pink) mice by incubation in Mg^2+^-free ACSF containing 6 mM KCl. The expanded recordings of the bursts (surrounded by grey rectangles) are shown on the right. Scale bars: 10 s (left)/200 ms (right), 50 µV. **G.** Quantification of burst frequency (top) and burst duration (bottom) in RACK1 fl/fl (white) and RACK1 cKO (pink) hippocampal slices (n=15 slices from 5 mice for RACK1 fl/fl, and n=18 slices from 6 mice for RACK1 cKO; unpaired t-test). **H.** Quantification of VU0134992’s effect on burst frequency (top) and duration (bottom) in RACK1 fl/fl (white) and RACK1 cKO (pink) hippocampal slices (RACK1 fl/fl: n=15 slices from 5 mice for burst frequency and duration, respectively; RACK1 cKO: n=18 slices from 6 mice for burst frequency and duration; repeated measures one-way ANOVA with Tukey’s multiple comparison test). The raw data are presented in **Table S2**.

We reasoned that the elevated expression of KIR4.1 K^+^ channels in RACK1 cKO astrocytes and the consequent enhancement in K^+^ buffering capacity might have major roles during intense neuronal activity. To test this hypothesis, we repeatedly stimulated SCs (10 Hz, 30 s) and analyzed the fEPSPs in CA1 region of the hippocampus. This stimulation induced rapid synaptic facilitation and then depression (**Fig. 7C**), which results from depletion of the presynaptic glutamate pool. Facilitation was greater and depression was slower in RACK1 cKO slices than in RACK1 fl/fl slices (**Fig. 7D**). This finding indicates that the elevated expression of KIR4.1 K^+^ channels in RACK1 cKO mice sustains repetitive excitatory synaptic activity. Accordingly, KIR4.1 K^+^ channel inhibition by VU0134992 (30 µM, for 20 min) had more of an effect on synaptic facilitation and depression in RACK1 cKO mice than in RACK1 fl/fl mice (**Fig. 7D**). Indeed, in the presence of VU0134992, repetitive stimulation-induced facilitation and subsequent depression were similar in RACK1 fl/fl and RACK1 cKO mice (**Fig. 7D**). These results indicate that astrocytic RACK1 regulates neuronal activity in response to repetitive stimulation by controlling the expression of KIR4.1 K^+^ channel.

We next tested the impact of RACK1 on recurrent burst activity. This was induced in hippocampal slices by incubation in a pro-epileptic artificial cerebrospinal fluid (ACSF) (Mg^2+^-free with 6 mM KCl (0Mg6K ACSF)). We recorded neuronal bursts in all hippocampal regions by using the multi-electrode array (MEA) technique (**Fig. 7E**). We found that bursts were less frequent and last for longer in RACK1 cKO mice than in RACK1 fl/fl mice (**Fig. 7F, G**). Hence, the burst rate under pro-epileptic conditions appeared to be better controlled in astrocytic RACK1 cKO mice than in RACK1 fl/fl mice. To check whether this was due to more efficient buffering of extracellular K^+^ released during sustained activity, we recorded burst activity in RACK1 fl/fl and RACK1 cKO slices in the presence of VU0134992 (30 µM). In RACK1 fl/fl mice, 15-25 min of inhibition of the KIR4.1 K^+^ channel by VU0134992 was associated with a transient increase in burst frequency that was likely due to the neuronal depolarization caused by extracellular K^+^ accumulation. This was followed (after >30 min of VU0134992 exposure) by a long-lasting decrease in burst frequency, which probably resulted from the accumulation of excess extracellular K^+^ (**Fig. 7H, top, left**). There was no effect on burst duration (**Fig. 7H, bottom, left**). Interestingly, this dual regulation of burst frequency was not observed in RACK1 cKO mice (**Fig. 7H, top, right**), which showed only a decrease in burst duration after >30 min treatment with VU0134992 (**Fig. 7H, bottom, right**). These results indicate that by controlling KIR4.1 expression and the associated K^+^ buffering capacity in astrocytes, RACK1 helps to modulate the firing rate when neuronal activity is sustained.

Collectively, our results show that RACK1 associates with specific mRNAs in astrocytes and, in particular, represses the translation of *Kcnj10* mRNA. This translational effect is mediated by the *Kcnj10* gene’s 5’UTR. RACK1 cKO in astrocytes is associated with higher KIR4.1 levels overall and in PAPs; in turn, this affects astrocyte volume and attenuates recurrent neuronal burst activity.

## Discussion

The objective of the present study was to investigate the molecular mechanisms that regulate translation in astrocytes. We developed a TRAP method for purifying polysome-associated proteins in astrocytes and focused on the 40S-associated protein RACK1, a critical factor in translational regulation (Gallo and Manfrini, 2015; Nielsen et al., 2017). We demonstrated that RACK1 interacts with specific mRNAs in astrocytes and PAPs, represses the translation of *Kcnj10* (encoding KIR4.1), and thus impacts astrocyte volume and neurotransmission.

The TRAP technique that we used to purify astrocyte polysomes was originally developed for the analysis of polysomal mRNAs (Doyle et al., 2008). Here, we demonstrated that TRAP was compatible with MS. As expected, the most abundant immunopurified proteins were ribosomal or translation complex-associated proteins, although other proteins were also identified. The most highly represented cytoskeletal associated proteins in our screen included Ckap4 (CLIMP63), an endoplasmic reticulum (ER) integral membrane protein that binds to microtubules and promotes ER tubule elongation (Vedrenne et al., 2005). Interestingly, Ckap4 has a crucial role in the dendritic organization of the ER in neurons (Cui-Wang et al., 2012). The cytoplasmic linker associated protein CLASP2 mediates asymmetric microtubule nucleation in the Golgi apparatus and is crucial for establishing the latter’s continuity and shape (Miller et al., 2009). CLASP2 cytoskeleton-related mechanisms have been shown to underlie microtubule stabilization, neuronal polarity and synapse formation and activity (Beffert et al., 2012). These proteins might be candidates for the regulation of translation in astrocytes. In contrast, our experiments did not pinpoint all the known RBPs in astrocytes; for instance, we did not detect Qki, which was recently shown to regulate translation in astrocytes (Radomska et al., 2013; Sakers et al., 2021). Thus, the TRAP-MS technique probably does not give a comprehensive view of the ribosome-associated proteome in the astrocyte. However, it is cell-specific and so might be a powerful approach for identifying some of the key post-transcriptional regulators in astrocytes.

Among the astrocytic ribosome-associated proteins identified in our screen, we focused on the highly enriched RACK1. Interestingly, use of a selection of astrocyte-specific mRNAs enabled us to determine the preferential association of RACK1 with *Kcnj10* and *Slc1a2* mRNAs. This finding indicated that RACK1-containing ribosomes associate with specific mRNAs in astrocytes and probably confer specific translational properties on the ribosomes. Along the same lines, it has been shown that ribosomes with different stoichiometries of RACK1 translate different subsets of mRNAs (Coyle et al., 2009). RACK1 was also recently described as one of the ribosomal proteins translated in neurites and able to rapidly go on and off the ribosomes in neurons – suggesting strongly that RACK1-containing ribosomes have specific functions (Fusco et al., 2021). Taken as a whole, these data suggest that RACK1 is involved in ribosome filtering mechanisms (Mauro and Matsuda, 2016). It remains to be seen how the interaction between RACK1-containing ribosomes and specific mRNAs is achieved but various elements might be involved in this process. RACK1 has been shown to discriminate between mRNAs according to their length and to promote the translation of mRNAs with a short open reading frame (Thompson et al., 2016). In viruses, RACK1 might mediate the translation of mRNAs with an internal ribosome entry site (Majzoub et al., 2014). Other studies have demonstrated that RACK1 controls translation by sensing 5’UTR sequences and structures (Gallo et al., 2018).

Here, we showed that cell-selective RACK1 KO led to higher levels of KIR4.1 in astrocytes and in PAPs, indicating that RACK1 represses *Kcnj10* translation. RACK1 has been shown to control important aspects of ribosome quality control by sensing stalled ribosomes on polyarginine or proline codons and contributing to the degradation of nascent protein chains on stalled ribosomes (Juszkiewicz et al., 2020; Kuroha et al., 2010; Sitron et al., 2017; Sundaramoorthy et al., 2017). We further confirmed this effect on a lysine AAA sequence. However, we showed that this type of mechanism does not operate on the *Kcnj10* CDS – indicating that the increase in KIR4.1 seen after RACK1 KO is not related to a ribosomal readthrough mechanism. In contrast, we showed that the RACK1-mediated control of *Kcnj10* relied on specific 5’UTR sequences. It now remains to be determined how this regulatory mechanism operates. With regard to RACK1’s mRNA selectivity, this question reamins extremely complex because several mechanisms might be involved. RACK1 recruits and controls the activity of translation factors like the elongation factor eIF6 (Rollins et al., 2019). RACK1 is known to interact with components of the microRNA-induced gene silencing complex. This interaction recruits the complex to the translation site and facilitates gene repression (Jannot et al., 2011). Baum et al. suggested that recruitment of the mRNA-binding protein Scp160 to the yeast homolog (Asc1p) of RACK1 may influence the translation of specific mRNAs (Baum et al., 2004). On the same lines, changes to the translation machinery recruited on the *Kcnj10* 5’UTR might occur in the absence of RACK1, which would change the ribosomal translational efficiency.

In previous research, we demonstrated that the RACK1-encoding gene *Gnb2l1* is preferentially translated in PAPs. This suggests that RACK1-ribosome composition in astrocytes is not homogenous and that RACK1 exerts its translational control preferentially in PAPs (Mazare et al., 2020b). We also determined that *Kcnj10* polysomal mRNAs are present in PAPs (Mazare et al., 2020b). In neuronal processes, the plasticity of ribosomal protein composition involves RACK1 (Fusco et al., 2021). Here, we found that RACK1 was associated with *Kcnj10* mRNAs in astrocytes in general but also in PAPs in particular. Moreover, we found that PAP levels of KIR4.1 were lower in the absence of RACK1. Taken as a whole, these data indicate that RACK1 regulates KIR4.1 translation not only in astrocytes in general but also locally in PAPs.

KIR4.1 is a weakly inwardly rectifying K^+^ channel; in astrocytes, it helps to maintain the resting membrane potential, high K^+^ conductance, volume regulation, and glutamate uptake (Chever et al., 2010; Djukic et al., 2007; Juszkiewicz and Hegde, 2017; Kucheryavykh et al., 2007; Olsen and Sontheimer, 2008; Seifert et al., 2009; Sibille et al., 2015). During neuronal activity, neurons release large amounts of K^+^ at the synapse. This K^+^ is rapidly taken up by astrocytic KIR4.1 and is then transported through the astrocytic network to regions with lower K^+^ levels. This astrocytic K^+^ clearance mechanism (known as “spatial K^+^ buffering” is vital for the maintenance of K^+^ homeostasis and the prevention of neuronal hyperexcitability. Defects in KIR4.1 expression or function are associated with various brain pathologies and with epilepsy in particular (Nwaobi et al., 2016). In mice, KIR4.1 inactivation or reduced expression of KIR4.1 in astrocytes affects neuronal function (as indicated by a reduction in hippocampal short-term plasticity) and leads to ataxic seizures and early death (Sibille et al., 2014).

Here, we showed that KIR4.1 overexpression did not modify basal excitatory synaptic transmission but was critical during sustained activity (10 Hz stimulation and recurrent bursts). Indeed, we observed greater facilitation and slower depression (compared with control mice) after repetitive stimulation in RACK1 cKO mice. This effect was abolished upon addition of a specific KIR4.1 blocker, thus demonstrating that the greater facilitation and slower depression observed in RACK1 cKO were related to the upregulation of KIR4.1. These results are consistent with previous reports of a role for KIR4.1 in the 3-10 Hz frequency band but not in the baseline activity (0.1 Hz) (Chever et al., 2010; Sibille et al., 2015). Regarding the neuronal network burst activity under pro-epileptic conditions (with 0Mg6K ACSF), the burst frequency was lower in RACK1 cKO than in RACK1 fl/fl mice. This finding suggests that the astrocytes were better able to buffer extracellular K^+^ through higher levels of KIR4.1, which enhanced the RACK1 cKO mice’s ability to control extracellular K^+^ levels and the firing rate. Taken as a whole, these data thus indicate that by modulating KIR4.1 expression, RACK1 regulates neurotransmission. Interestingly, a relationship between RACK1 and epilepsy has been reported previously, albeit without a focus on astrocytes. RACK1 was shown to repress the transcription of the voltage-gated sodium channel α subunit type I (SCN1A) mRNA, the downregulation of which is associated with epilepsy (Dong et al., 2014). In the lithium-pilocarpine rat model, RACK1 levels in the hippocampus are elevated after epileptic episodes (Xu et al., 2015). Furthermore, in a rat model of mesial temporal lobe epilepsy, RACK1 was upregulated in the granular layer dorsal dentate gyrus and downregulated in the ventral dentate gyrus (do Canto et al., 2020). These literature data and our present results suggest that RACK1 is an important factor in the regulation of neurotransmission.

The present study focused on mechanisms of translation in astrocytes. We found that in astrocytes, RACK1 is a ribosomal associated protein able to interact selectively with mRNAs. We showed that RACK1 represses the synthesis of KIR4.1, which has a critical role in maintaining extracellular K^+^ levels in the brain. Dysfunction of KIR4.1 in rodents and humans evokes seizures and chronic epilepsy. Thus, through the regulation of KIR4.1 levels, RACK1 might be a therapeutic target in epilepsy.

## Supporting information

Key resource table

## Acknowledgements

We are grateful to the donors who support the charities and charitable foundations cited below. This work was funded by a grant from the *Fondation pour la Recherche Médicale* (FRM, grant reference AJE20171039094) to M. C.-S., and by the European Research Council (Consolidator grant #683154) to N.R.. The creation of the Center for Interdisciplinary Research in Biology (CIRB) was funded by the Fondation Bettencourt Schueller. We thank Robin Rondon and Ines Masurel for their help with RNA immunoprecipitation experiments. Lastly, we thank Julien Dumont and Philippe Mailly (Orion CIRB imaging platform) for help with imaging and analysis. The pSpCas9(BB)-2A-Puro vector (Addgene plasmid 62988) was a gift from Feng Zhang (https://zlab.bio/), and pmGFP-P2A-K0-P2A-RFP and pmGFP-P2A-K(AAA)20-P2A-RFP (Addgene plasmids 105686 and 105688) were gifts from Ramanujan Hegde (MRC Laboratory of Molecular Biology). The flow cytometry experiments were performed at the Imagerie-Gif core facility, which is funded by the French Agence Nationale de la Recherche (ANR-11-EQPX-0029/Morphoscope, ANR-10-INBS-04/FranceBioImaging; ANR-11-IDEX-0003-02/ Saclay Plant Sciences). Copy-editing assistance was provided by Biotech Communication SARL (Ploudalmézeau, France).

## Methods

### Animal care and ethical approval

Tg (Aldh1l1-eGFP/Rpl10a) JD130Htz (MGI: 5496674) (bacTRAP, **BT**) mice were obtained from Nathaniel Heintz’s laboratory (Rockefeller University, New York City, NY) and kept under pathogen-free conditions (Heiman et al., 2014). The genotyping protocol is described on the bacTRAP project’s website (www.bactrap.org). Tg(Aldh1l1-cre/ERT2)1Khakh (MGI:5806568) (Aldh1l1-Cre/ERT2) mice (Srinivasan et al., 2016) were obtained from the Jackson laboratory (https://www.jax.org/) and B6J.Cg-Rack1^tm1.1Cart^/Mmucd (RACK1 fl/fl) (MMRRC 044021-UCD) from the mutant mouse resource and research center (MMRRC) (https://www.mmrrc.org/)(Cheng and Cartwright, 2018). C57BL6 WT mice were purchased from Janvier Labs (Le Genest-Saint-Isle, France). Mice were maintained on a C57BL6 genetic background. All experiments were performed on 2-month-old mice. Both sexes were used for all experiments.

Mice were kept in pathogen-free conditions. All animal experiments were carried out in compliance with (i) the European Directive 2010/63/EU on the protection of animals used for scientific purposes and (ii) the guidelines issued by the French National Animal Care and Use Committee (reference: 2013/118). The study was also approved by the French Ministry for Research and Higher Education’s institutional review board (reference 21817).

### Tamoxifen induction of RACK1 inactivation

Two-month-old mice received a daily intraperitoneal injection of 100 mg/kg tamoxifen solution in corn oil (10 mg/ml dissolved extemporaneously for 6-8h at 37°C) for 5 consecutive days and were analyzed 3 weeks later. For controls, RACK1 fl/fl received corn oil only (immunofluorescence; Western blot; qPCR), or tamoxifen (astrocyte volume study; electrophysiology)

### TRAP-MS

Whole brain homogenates (one brain per sample) from 2-month-old C57Bl/6 mice (WT, negative control) and BT mice were submitted to TRAP by immunoprecipitating GFP-fused astrocytic polyribosomes with anti-GFP antibodies and protein-G-coupled magnetic beads, as described elsewhere (Mazare et al., 2020a), except that 1 mg of proteins were used for the immunoprecipitation on 25 𝜇L G-protein-coupled magnetic Dynabeads coated with anti-GFP antibodies at 4°C. At the end of the procedure, immunoprecipitated proteins were eluted by boiling the beads in 20 µL of 0.35 M KCl buffer with 5X Laemmli buffer for 5 min. Samples were run on SDS-PAGE gels (Invitrogen) without separation as a clean-up step and then stained with colloidal blue staining (LabSafe GEL Blue^TM^ G Biosciences). Gel slices were excised, and proteins were reduced with 10 mM DTT prior to alkylation with 55 mM iodoacetamide. After washing and shrinking the gel pieces with 100% acetonitrile, in-gel digestion was performed using 0.10 µg trypsin/Lys-C (Promega) overnight in 25 mM NH_4_HCO_3_ at 30 °C. Peptides were then extracted (using 60/35/5 acetonitrile/H_2_O/HCOOH) and vacuum concentrated to dryness. Peptides were reconstituted in injection buffer (0.3% TFA) before LC-MS/MS analysis. Five replicates per conditions were prepared.

#### LC-MS/MS analysis

Online chromatography was performed with an RSLCnano system (Ultimate 3000, Thermo Scientific) coupled to a Q Exactive HF-X. Peptides were first trapped onto a C18 column (75 µm inner diameter × 2 cm; nanoViper Acclaim PepMapTM 100, Thermo Scientific) with buffer A (0.1% formic acid) at a flow rate of 2.5 µL/min over 4 min. The peptides were separated on a 50 cm x 75 µm C18 column (nanoViper C18, 3 µm, 100 Å, Acclaim PepMap^TM^ RSLC, Thermo Scientific) at 50°C, with a linear gradient from 2% to 30% buffer B (100% acetonitrile, 0.1% formic acid) at a flow rate of 300 nL/min over 91 min. Full MS scans were performed with the ultrahigh-field Orbitrap mass analyzer in the range m/z 375–1500, with a resolution of 120,000 at m/z 200. The top 20 intense ions were subjected to Orbitrap for further fragmentation via high energy collision dissociation activation and a resolution of 15,000, with the intensity threshold kept at 1.3 × 10^5^. We selected ions with a charge from 2+ to 6+ for screening. The normalized collision energy was set to 27 and the dynamic exclusion was set to 40 s.

#### Data analysis

Data were searched against the *Mus musculus* UniProt canonical database (downloaded in August 2017 and containing 16888 sequences) using Sequest HT via proteome discoverer (version 2.0). The enzyme specificity was set to trypsin, and a maximum of two missed cleavage sites was allowed. Oxidized methionine, carbamidomethyled cysteine, and N-terminal acetylation were set as variable modifications. The maximum allowed mass deviation was set to 10 ppm for monoisotopic precursor ions and to 0.02 Da for MS/MS peaks. The resulting files were further processed using myProMS (Poullet et al., 2007) version 3.9.3 (https://github.com/bioinfo-pf-curie/myproms). The false-discovery rate (FDR) was calculated using Percolator (The et al., 2016) and was set to 1% at the peptide level for the whole study. Label-free quantification was performed using peptide extracted ion chromatograms (XICs) computed with MassChroQ (Valot et al., 2011) v.2.2.1. For protein quantification, XICs from proteotypic peptides shared between compared conditions (TopN matching) with two-missed cleavages were used. Median and scale normalization at the peptide level was applied to the total signal, in order to correct the XICs in each biological replicate. To estimate the significance of the change in protein abundance, a linear model (adjusted for peptides and biological replicates) was used, and p-values were adjusted using the Benjamini–Hochberg FDR procedure. Proteins with at least three total peptides in all replicates, a two-fold enrichment and an adjusted p-value ≤ 0.05 were considered to be significantly enriched in sample comparisons. Proteins only found in one condition were also considered if they matched the peptide criteria. Proteins selected with these criteria were further analyzed and subjected to a GO functional enrichment analysis. The raw MS proteomics data have been deposited to the ProteomeXchange Consortium via the PRIDE (Perez-Riverol et al., 2019) partner repository (http://www.ebi.ac.uk/pride), with the dataset identifier PXD033121.

### GO analysis

A GO analysis was performed for proteins with at least three peptides read by LC-MS/MS and found to be enriched in BT extracts (p-value < 0.05 and Log2 FC > 1), using UniProt bank annotations for the mouse (UniProt-GOA Mouse - *Mus musculus*). GO-term-associated p-values were computed with the GOTermFinder module of myProMS (Poullet et al., 2007). We analyzed biological processes and molecular functions (p-value threshold: 0.05). For each family, GO terms were classified manually according to the GO hierarchy, taking into account the number of genes from the study included in the highest GO. For instance, the in the “Gene Expression” category were included in the highest GO “Metabolic process”, and the proteins in the ‘Ribosomal proteins’ category were included in “Structural molecule activity”. The number of proteins in each category was expressed as a percentage of the total number of proteins. These data should be taken as illustrative because some proteins have more than one role and so the categories overlap.

### RACK1 immunoprecipitation (IP)

IP was performed according to the TRAP-MS protocol (i.e. using an anti-RACK1 antibody) but with some changes, as follows. Columns (bead volumes: 100𝜇L for the precleaning column, 25𝜇L for precleaning + IgG column, 25 𝜇L for IP column) were prepared the day before. The IP column was first blocked 1 h with 2% bovine serum albumin and 0.1 mg/100 µL beads of yeast tRNA in 0.15 M KCl buffer, rinsed with 0.15 M KCl three times and coated with 5 𝜇g of anti-RACK1 antibodies or 5 𝜇g of non-specific immunoglobulins IgG (negative control). 500 𝜇g of protein extract was used. The precleaning steps have been described elsewhere (Mazare et al., 2020a). The precleaned extract was incubated with IP columns for 30 min at 4°C. The beads were rinsed three times with 0.35 M KCl and RNA were eluted in 300 𝜇L RLT buffer (Qiagen, Hilden, Germany) for 5 min at room temperature (RT) and kept at −80°C until extraction.

### Quantitative RT-PCR

RNA was extracted using the Rneasy kit (Qiagen, Hilden, Germany). cDNA was then generated using the Superscript™ III Reverse Transcriptase kit (ThermoFisher). Differential levels of cDNA expression were measured using the droplet digital PCR (ddPCR) system (Bio-Rad) and TaqMan® copy number assay probes or primers (**Key resource table**). Briefly, cDNA and 6-carboxyfluorescein probes or primers were distributed into 10,000–20,000 droplets. The nucleic acids were then PCR-amplified in a thermal cycler and read (as the number of positive and negative droplets) with a QX200 ddPCR system. The results were normalized as follows: the IgG IP results were subtracted from the RACK1 RNA IP results for each gene. The results were then normalized against 18S rRNA gene expression. For GFP RNA IP (TRAP), results were normalized against the 18S rRNA.

### Brain slices for FISH and immunofluorescence

Mice were anesthetized with a mix of ketamine/xylazine (80/100 mg/kg i.p.) and killed by transcardiac perfusion with PBS/PFA 4%. The brain was removed, incubated in 30% sucrose overnight, and cut into 40-µm-thick sections using a cryomicrotome (HM 450, Thermo Scientific). For long-term storage, slices were kept at −20°C in a cryoprotectant solution (30% ethylene glycol, 30% glycerol, 40% PBS).

### High-resolution FISH and GFAP co-immunofluorescent detection and analysis

FISH was performed using the v2 Multiplex RNAscope technique (Advanced Cell Diagnostics, Inc., Newark, CA, USA). After the FISH procedure, GFAP was detected via immunofluorescence. Astrocyte-specific FISH dots were identified from their position on the GFAP immunolabeling image, using the *AstroDot* ImageJ plug-in. This method has been described in detail elsewhere (Oudart et al., 2020).

### Immunohistochemical labeling and confocal imaging

Immunohistochemical labeling was performed on frozen brain sections (see above) rinsed in PBS and incubated for 2 h at RT in blocking solution (5% normal goat serum, 0.5% Triton X-100 in PBS). Sections were incubated with primary antibodies diluted in the blocking solution overnight at 4°C, rinsed for 5 min in PBS three times, incubated with secondary antibodies diluted in blocking solution for 2 h at RT, rinsed for 5 min in PBS three times, and mounted in Fluoromount (Southern Biotech, Birmingham, AL). Brain sections were imaged on X1 or W1 spinning-disk confocal microscopes (Yokogawa). Images were acquired with a 40X oil immersion objective (Zeiss). For the astrocyte morphology study, a LSM 980 confocal (Zeiss) and a 63X oil immersion objective (Zeiss) were used. The antibodies used in the present study are listed in **Key resource table**.

### Preparation of synaptogliosomes

Synaptosomes were prepared as described elsewhere (Mazare et al., 2020a). All steps were performed at 4°C. Hippocampi (two per extract; 1 mouse) were dissected and homogenized with a tight glass homogenizer (20 strokes) in buffer solution (0.32 M sucrose and 10 mM HEPES in DNAse/RNAse-free water, with 0.5 mM DTT, protease inhibitors (cOmplete^TM^, EDTA free, 1 minitablet/10 mL), ribonuclease inhibitor (1 µL/mL, cycloheximide (CHX) 100 µg/mL freshly prepared). The homogenate was centrifuged at 900 g for 15 min. The pellet was discarded, and the supernatant was centrifuged at 16,000 g for 15 min. The new supernatant was discarded, and the pellet (containing synaptogliosomes) was diluted in 600 µl of buffer solution and centrifuged again at 16,000 g for 15 min. The final pellet contained the synaptogliosomes.

### Western blots

Whole hippocampi were crushed with a pestle and a mortar at −80°C. Proteins were extracted from the tissue powder or synaptogliosome pellets in 2% SDS (500 µl or 200 µl per sample, respectively) with EDTA-free Complete Protease Inhibitor (Roche), sonicated twice for 5 min or once for 5 min, respectively (Bioruptor UCD 200, diagenode), and centrifuged at 20,000 g for 20 min at 4°C. Supernatants were heated at 56°C in Laemmli loading buffer for 5 min. Protein content was measured using the Pierce 660 nm protein assay reagent (Thermo Scientific) and the Multiskan™ FC spectrophotometer (Thermo Scientific). Equal amounts of protein (whole immunoprecipitation extracts: 10 to 20 µg for hippocampus and synaptogliosomes) were separated by denaturing electrophoresis in Mini-Protean TGX stain-free gels (Biorad) and then electrotransferred to nitrocellulose membranes using the Trans-blot Turbo Transfer System (Biorad). Membranes were hybridized as described previously (Ezan et al., 2012). The antibodies used in this study are listed in **Key resource table**. Horseradish peroxidase activity was visualized by enhanced chemiluminescence in a Western Lightning Plus system (Perkin Elmer, Waltham, MA, USA). Chemiluminescent imaging was performed on a Fusion FX system (Vilber). The chemiluminescence signal intensity for each antibody was normalized against that of stain-free membranes.

### Representative structure of the human 80S ribosome

The human 80S ribosome’s representative structure was depicted using the PyMol software (version 2.3.4, python 3.7, https://pymol.org/2/). A high-resolution cryo-electron microscopy (EM) structure of the human 80S ribosome (Natchiar et. al., 2017) was obtained from the Protein Data Bank in Europe (code 6EK0) because the mouse 80S ribosome is not available. The chain codes can be found on the Protein Data Bank in Europe website. The Lz chain for RPL10a and the Sg chain for RACK1 were selected.

### Cell lines and culture conditions

HEK293T (Thermo Fisher Scientific, Waltham, MA) cells were cultured in DMEM supplemented with 10% fetal bovine serum (FBS) and 1% penicillin/streptomycin (P/S) (Wisent Technologies). Control and RACK1 KO HEK293 cells were maintained in DMEM supplemented with 10% FBS, 1% P/S, 100 µg/mL zeocin (Thermo Fisher Scientific, R25001), and 15 µg/mL blasticidin (Thermo Fisher Scientific, R210-01). All cells were cultured at 37°C, in a humidified atmosphere with 5% CO_2_.

### Plasmid constructs

To generate the *Kcnj10* CDS containing a fluorescent reporter, a control cassette was first created by replacing the BspEI/KpnI segment of the pmGFP-P2A-K0-P2A-RFP (Addgene plasmids 105686) with a linker containing a P2A site, a Flag coding sequence, and the EcoRI and NotI restriction sites. A gene block (Integrated DNA Technologies) encoding mKIR4.1 (AAI41089.1) without a stop codon was then inserted at the EcoRI/NotI sites of this control cassette in frame with both the GFP and mCherry coding sequences. The psiCHECK-2 vector (Promega, C8021) was used to build the dual luciferase reporters with *Kcnj10* UTRs. The UTR sequences of mouse *Kcnj10* mRNA (NM_001039484.1 and AB039879.1) were synthesized as gBlocks and inserted at the NheI site of the psiCHECK-2 vector. The 3’UTR of the *Kcnj10* mRNA (AB039879.1) was inserted as a gBlock into the XhoI and NotI restriction sites in the psiCHECK-2 vector downstream of the *Renilla* luciferase reporter gene. The truncated versions of the *Kcnj10* 5′ UTR (1-146; 127-242; 95-242; 95-191; 1-191; 181-242) were inserted as NheI/NheI PCR fragments into the psiCHECK-2 vector at the 5′ end of the *Renilla* luciferase gene. The sequences of the primers and gBlocks used for subcloning are listed in **Key resource table**.

### Flow cytometry analysis

Transient transfection of fluorescent reporter constructs was performed using Lipofectamine 2000 (Thermo Fisher Scientific), according to manufacturer’s instructions. In all experiments, WT and RACK1^KO^ HEK293T cells were plated in 6-well plates at a concentration of 500,000 cells per well, and transfected with 10 ng of plasmids on the following day. The cells were then trypsinized, washed once with PBS and pelleted at RT at 500 g for 5 min. The cells were resuspended in 500 µl PBS containing 10% FBS, passed through a 40 µm filter, and analyzed with a CytoFlex flow cytometer (Beckman Coulter). 10,000 fluorescent cells were selected for the analysis of GFP and mCherry signals. The data were analyzed using FlowJo software.

### CRISPR/cas9-mediated genome editing

CRISPR-Cas9-mediated genome editing of HEK293 cells was performed according to the method described by Ran *et al*. (Ran et al., 2013). The DNA oligonucleotides (encoding a small guide RNAs (sgRNAs) cognate to the coding region of human *Rack1*/*Gnb2l1* gene) are detailed in **Key resource table**. These oligos contained BbsI restriction sites and were annealed to create overhangs for cloning of the guide sequence oligos into pSpCas9(BB)-2A-Puro (PX459) V2.0 (Addgene plasmid 62988) by BbsI digestion. To generate KO HEK293T cells, we transfected 500,000 cells with the guide sequence containing the pSpCas9(BB)−2A-puro plasmid. Twenty-four hours after transfection, puromycin was added to the cell medium. After 72 h, puromycin-resistant cells were isolated in 96-well plates and cultured until monoclonal colonies were obtained. Clonal cell populations were analyzed for protein depletion in Western blots.

### Dual luciferase reporter assays

WT or RACK1^KO^ HEK293T cells were transfected with 20 ng per well of each psiCHECK2 construct or the empty psiCHECK2 in a 24-well plate by using Lipofectamine 2000 (Thermo Scientific, 11668019), according to the manufacturer’s instructions. Cells were lysed 24 h after transfection, and luciferase activities were measured with the Dual-Luciferase Reporter Assay System (Promega) in a GloMax 20/20 luminometer (Promega). The RLuc activity was normalized against the activity of co-expressed FLuc, and the normalized RLuc values were quoted relative to the corresponding control.

### Viral vectors and stereotaxic injection

Two-month-old mice were anesthetized with a mixture of ketamine (95 mg/kg; Merial) and xylazine (10 mg/kg; Bayer) in 0.9% NaCl and placed on a stereotaxic frame with constant body temperature monitoring. AAVs were diluted in PBS with 0.01% Pluronic F-68 at a concentration of 9×10^12^ vg/ml and 1 µl of virus was injected bilaterally into the hippocampus at a rate of 0.1 µl/min, using a 29-gauge blunt-tip needle linked to a 2 µl Hamilton syringe (Phymep). The stereotaxic coordinates relative to the bregma were as follows: anteroposterior, ±2 mm; mediolateral: +1.5 mm; dorsoventral, − 1.5 mm. The needle was left in place for 5 min and then removed slowly. The skin was glued back in place, and the animals’ recovery was checked regularly for the next 24 h. After 11 days, the mice were sacrificed and the tissues were processed for immunofluorescence assays.

### Measurement of astrocyte volume

To drive expression in astrocytes, the transgene encoding cytosolic red fluorescent protein Td tomato was inserted under the control of the gfaABC1D (Lee et al., 2008) into an AAV shuttle plasmid containing the inverted terminal repeats of AAV2. Pseudotyped serotype 9 AAV particles were produced by transient co-transfection of HEK-293T cells, as described previously (Fol et al., 2016). Viral titers were determined by quantitative PCR amplification of the inverted terminal repeats on DNase-resistant particles and were expressed in vg per ml.

Astrocytes on 100 µm brain sections were reconstructed in 3D, using IMARIS software (Oxford Instruments, version 9.7.2). Filaments were created with a unique starting point in the astrocyte soma and with seeds defined with a manual threshold, according to the fluorescence intensity. Filaments outside the astrocyte were removed manually. An envelope of the astrocyte territory was created using the convex hull plugin (Matlab). The following variables were computed and exported for analysis: astrocyte volume (corresponding to the envelope volume), the sum of the filament length and data for a 3D Sholl analysis (5 μm steps).

### Electrophysiology

Electrophysiological recordings were performed in the hippocampus of 3-month-old RACK1 fl/fl (Control) and RACK1 cKO mice 3 weeks after tamoxifen injection, using ACSF in the presence or absence of a Kir4.1 blocker (30 µM VU0134992, Tocris, Biotechne) (Kharade et al., 2018).

#### Acute hippocampal slice preparation

Acute transverse hippocampal slices (400 µm) were prepared as described previously (Chever et al., 2016) from 3-month-old RACK1 fl/fl or astrocytic RACK1 cKO mice. Briefly, slices were cut at low speed (0.04 mm/s) and at a vibration frequency of 70 Hz in ice-cold oxygenated ACSF supplemented with sucrose (in mM: 87 NaCl, 2.5 KCl, 2.5 CaCl_2_, 7 MgCl_2_, 1 NaH_2_PO_4_, 25 NaHCO_3_ and 10 glucose, saturated with 95% O_2_ and 5% CO_2_). Slices were then maintained at 32°C in a storage chamber containing standard ACSF (in mM: 119 NaCl, 2.5 KCl, 2.5 CaCl_2_, 1.3 MgSO_4_, 1 NaH2PO_4_, 26.2 NaHCO_3_ and 11 glucose, saturated with 95% O_2_ and 5% CO_2_), for at least 1 h prior to recording.

#### Field recordings

Slices were transferred to a submerged recording chamber mounted on an Olympus BX51WI microscope equipped for infrared-differential interference microscopy and were perfused with standard ACSF at a rate of 1-2 ml/min at 32°C. Extracellular field recordings were performed with glass pipettes (2–5 MΩ) filled with ACSF and placed in the stratum radiatum. Stimulus artifacts were blanked in sample recordings. Basal excitatory synaptic transmission (input/output curves) was evaluated in presence of picrotoxin (100 µM), and the tissue was cut between CA1 and CA3 to prevent the propagation of epileptiform activity. Evoked postsynaptic responses were induced by stimulating SCs at 0.1 Hz in the CA1 *stratum radiatum*. Slices underwent prolonged, repetitive stimulation at 10 Hz for 30 s. Responses (neuronal fEPSP slope) were binned (bin size: 1.2 s) and normalized against the mean baseline response measured at 0.1 Hz prior to repetitive stimulation. Both basal excitatory synaptic transmission and responses to repetitive stimulation were evaluated before and after treatment with VU0134992. Field potentials were acquired with Axopatch-1D amplifiers (Molecular Devices), digitized at 10 kHz, filtered at 2 kHz, and stored and analyzed on a computer using pCLAMP9 and Clampfit10 software (Molecular Devices).

#### MEA recordings

MEA recordings were performed as described previously (Chever et al., 2016). After a 20 min incubation in standard ACSF at 32°C, slices were stored for at least 1 h before recording in magnesium-free ACSF containing 6 mM KCl (0Mg6K ACSF) at 32°C. Hippocampal slices were then transferred onto planar MEA petri dishes (200-30 indium tin oxide electrodes, organized in a 12×12 matrix, with an internal reference, 30 µm diameter and 200 µm inter-electrode distance; Multichannel Systems), kept in place with a small platinum anchor, and continuously perfused at 1-2 ml/min with 0Mg6K ACSF at 32°C. Pictures of cortical slices on MEAs were acquired with a video microscope table (MEA-VMT1; Multichannel Systems). MEA_Monitor software (Multichannel Systems) was used to identify the location of the electrodes relative to the various regions of the hippocampal. Data were sampled at 10 kHz, and the slice activity was recorded at 32°C using a MEA2100-120 system (bandwidth: 1-3000 Hz; gain: 5x; Multichannel Systems) and MC_Rack 4.5.1 software (Multichannel Systems). The slices’ activity was recorded in 0Mg6K ACSF before and after treatment with 30 µM VU0134992. Raw data on 0Mg6K ACSF-induced network burst activity was analyzed with MC Rack software (Multichannel Systems). Bursts were detected with the Spike Sorter algorithm, which sets a threshold based on multiples of the standard deviation of the noise calculated over the first 500 ms of recording free of electrical activity. A 5-fold standard deviation threshold was used to automatically detect each event. If required (after a visual check), each event could be modified in real-time by the operator. Bursts were defined arbitrarily as discharges lasting less than 5 s. The bursts were characterized by fast voltage oscillations and then slow oscillations or negative shifts. The burst duration was measured using Neuroexplorer software (version 4.109, Nex Technologies, USA).

### Statistics

All statistical analyses were performed using GraphPad Prism software (version 8.0.2, GraphPad Software, Inc.). The statistical tests are listed in Table S2 and in the figure legends. For the analysis of qPCR and Western blot data, a t-test was applied if the data were normally distributed (according to the Kolmogorov-Smirnov test) and the variances were equal (according to Fisher’s test); if not, a Mann-Whitney test was applied. For *in cellulo* studies, a t-test was used with Welch’s correction if the variances were equal; if not, a Mann-Whitney test was used.

### Supplementary Informations

**Figure S1: RACK1’s stalling effect on a reporter sequence**

**A.** Schematic representation of the reporter constructs used to quantify ribosome stalling. The reporter contains GFP and mCherry separated by a sequence either lacking or containing 20 AAA lysine codons (K_0_ and K_20_, respectively) and surrounded by viral 2A sequences. **B.** A representative experiment monitoring the fluorescence protein ratio in WT or RACK1^KO^ HEK293T cells transfected with the K_0_ and K_20_ reporters. **C.** A histogram of the data, presented as the mean ± SD (N = 3); an unpaired MannWhitney test and a two-tailed t-test with Welch’s correction. ns, not significant, p-value>0.05; ***, p<0.001. The raw data are presented in Table S2.

**Figure S2: Alignment of the 5’UTR sequences of *Kcnj10***

**A.** The sequences were aligned using Clustal Omega and default parameters, and colored in Jalview by identity. **B.** Plot showing the GC content in the 5’UTR sequence of murine *Kcnj10* mRNA. The schematic representation shows two GC-rich regions (blue boxes).

**Figure S3: RACK1’s sensitivity to Kcnj10 5’UTR is not mediated by an effect on mRNA levels**

**A.** Schematic representation of the *Renilla* luciferase (RLuc) reporter constructs harboring the 5’UTR of mouse *Kcnj10* mRNA. These sequences were inserted in the psiCHECK2 vector, which also encodes the Firefly luciferase (Fluc). The “control” RLuc was generated by transfecting the empty psiCHECK2 vector. **B.** The qPCR ratio of *Renilla* luciferase (RLuc) to firefly luciferase (Fluc) mRNA levels in WT and RACK1^KO^ HEK293T cells transfected with constructs empty of harboring the 5’UTR #1 or 2 of mouse *Kcnj10* mRNA.

**Table S1: The raw data for TRAP-MS and GO analyses**

**1.** Parameters used for the LC-MS-MS study. **2.** Comparison of immunoprecipitation data in WT vs. BT extracts. **3.** FC, fold change. GO analysis of biological processes. **4.** GO analysis of molecular functions

**Table S2: Datasets**

## References

Battaini, F., and Pascale, A. (2005). Protein kinase C signal transduction regulation in physiological and pathological aging. Ann N Y Acad Sci 1057, 177–192.

Battaini, F., Pascale, A., Lucchi, L., Pasinetti, G.M., and Govoni, S. (1999). Protein kinase C anchoring deficit in postmortem brains of Alzheimer’s disease patients. Exp Neurol 159, 559–564.

Baum, S., Bittins, M., Frey, S., and Seedorf, M. (2004). Asc1p, a WD40-domain containing adaptor protein, is required for the interaction of the RNA-binding protein Scp160p with polysomes. Biochem J 380, 823–830.

Beffert, U., Dillon, G.M., Sullivan, J.M., Stuart, C.E., Gilbert, J.P., Kambouris, J.A., and Ho, A. (2012). Microtubule plus-end tracking protein CLASP2 regulates neuronal polarity and synaptic function. J Neurosci 32, 13906–13916.

Besse, F., and Ephrussi, A. (2008). Translational control of localized mRNAs: restricting protein synthesis in space and time. Nat Rev Mol Cell Biol 9, 971–980.

Blanco-Urrejola, M., Gaminde-Blasco, A., Gamarra, M., de la Cruz, A., Vecino, E., Alberdi, E., and Baleriola, J. (2021). RNA Localization and Local Translation in Glia in Neurological and Neurodegenerative Diseases: Lessons from Neurons. Cells 10.

Boulay, A.C., Saubaméa, B., Adam, N., Chasseigneaux, S., Mazaré, N., Gilbert, A., Bahin, M., Bastianelli, L., Blugeon, C., Perrin, S., et al. (2017). Translation in astrocyte distal processes sets molecular heterogeneity at the gliovascular interface. Cell Discovery 3, 17005.

Carney, K.E., Milanese, M., van Nierop, P., Li, K.W., Oliet, S.H., Smit, A.B., Bonanno, G., and Verheijen, M.H. (2014). Proteomic analysis of gliosomes from mouse brain: identification and investigation of glial membrane proteins. J Proteome Res 13, 5918–5927.

Cheng, Z.F., and Cartwright, C.A. (2018). Rack1 maintains intestinal homeostasis by protecting the integrity of the epithelial barrier. Am J Physiol Gastrointest Liver Physiol 314, G263–G274.

Chever, O., Djukic, B., McCarthy, K.D., and Amzica, F. (2010). Implication of Kir4.1 channel in excess potassium clearance: an in vivo study on anesthetized glial-conditional Kir4.1 knock-out mice. J Neurosci 30, 15769–15777.

Chever, O., Dossi, E., Pannasch, U., Derangeon, M., and Rouach, N. (2016). Astroglial networks promote neuronal coordination. Sci Signal 9, ra6.

Cohen-Salmon, M., Slaoui, L., Mazare, N., Gilbert, A., Oudart, M., Alvear-Perez, R., Elorza-Vidal, X., Chever, O., and Boulay, A.C. (2021). Astrocytes in the regulation of cerebrovascular functions. Glia 69, 817–841.

Coyle, S.M., Gilbert, W.V., and Doudna, J.A. (2009). Direct link between RACK1 function and localization at the ribosome in vivo. Mol Cell Biol 29, 1626–1634.

Cui-Wang, T., Hanus, C., Cui, T., Helton, T., Bourne, J., Watson, D., Harris, K.M., and Ehlers, M.D. (2012). Local zones of endoplasmic reticulum complexity confine cargo in neuronal dendrites. Cell 148, 309–321.

Culver, B.P., Savas, J.N., Park, S.K., Choi, J.H., Zheng, S., Zeitlin, S.O., Yates, J.R., 3rd, and Tanese, N. (2012). Proteomic analysis of wild-type and mutant huntingtin-associated proteins in mouse brains identifies unique interactions and involvement in protein synthesis. J Biol Chem 287, 21599–21614.

Dallerac, G., Zapata, J., and Rouach, N. (2018). Versatile control of synaptic circuits by astrocytes: where, when and how? Nat Rev Neurosci 19, 729–743.

Djukic, B., Casper, K.B., Philpot, B.D., Chin, L.S., and McCarthy, K.D. (2007). Conditional knock-out of Kir4.1 leads to glial membrane depolarization, inhibition of potassium and glutamate uptake, and enhanced short-term synaptic potentiation. J Neurosci 27, 11354–11365.

do Canto, A.M., Vieira, A.S., A, H.B.M., Carvalho, B.S., Henning, B., Norwood, B.A., Bauer, S., Rosenow, F., Gilioli, R., Cendes, F., et al. (2020). Laser microdissection-based microproteomics of the hippocampus of a rat epilepsy model reveals regional differences in protein abundances. Sci Rep 10, 4412.

Dong, Z.F., Tang, L.J., Deng, G.F., Zeng, T., Liu, S.J., Wan, R.P., Liu, T., Zhao, Q.H., Yi, Y.H., Liao, W.P., et al. (2014). Transcription of the human sodium channel SCN1A gene is repressed by a scaffolding protein RACK1. Mol Neurobiol 50, 438–448.

Doyle, J.P., Dougherty, J.D., Heiman, M., Schmidt, E.F., Stevens, T.R., Ma, G., Bupp, S., Shrestha, P., Shah, R.D., Doughty, M.L., et al. (2008). Application of a translational profiling approach for the comparative analysis of CNS cell types. Cell 135, 749–762.

Ezan, P., Andre, P., Cisternino, S., Saubamea, B., Boulay, A.C., Doutremer, S., Thomas, M.A., Quenech’du, N., Giaume, C., and Cohen-Salmon, M. (2012). Deletion of astroglial connexins weakens the blood-brain barrier. J Cereb Blood Flow Metab 32, 1457–1467.

Fol, R., Braudeau, J., Ludewig, S., Abel, T., Weyer, S.W., Roederer, J.P., Brod, F., Audrain, M., Bemelmans, A.P., Buchholz, C.J., et al. (2016). Viral gene transfer of APPsalpha rescues synaptic failure in an Alzheimer’s disease mouse model. Acta Neuropathol 131, 247–266.

Fusco, C.M., Desch, K., Dorrbaum, A.R., Wang, M., Staab, A., Chan, I.C.W., Vail, E., Villeri, V., Langer, J.D., and Schuman, E.M. (2021). Neuronal ribosomes exhibit dynamic and context-dependent exchange of ribosomal proteins. Nat Commun 12, 6127.

Gallo, S., and Manfrini, N. (2015). Working hard at the nexus between cell signaling and the ribosomal machinery: An insight into the roles of RACK1 in translational regulation. Translation (Austin) 3, e1120382.

Gallo, S., Ricciardi, S., Manfrini, N., Pesce, E., Oliveto, S., Calamita, P., Mancino, M., Maffioli, E., Moro, M., Crosti, M., et al. (2018). RACK1 Specifically Regulates Translation through Its Binding to Ribosomes. Mol Cell Biol 38.

Gay, D.M., Lund, A.H., and Jansson, M.D. (2022). Translational control through ribosome heterogeneity and functional specialization. Trends Biochem Sci 47, 66–81.

Harvey, R.F., Smith, T.S., Mulroney, T., Queiroz, R.M.L., Pizzinga, M., Dezi, V., Villenueva, E., Ramakrishna, M., Lilley, K.S., and Willis, A.E. (2018). Trans-acting translational regulatory RNA binding proteins. Wiley Interdiscip Rev RNA 9, e1465.

Heiman, M., Kulicke, R., Fenster, R.J., Greengard, P., and Heintz, N. (2014). Cell type-specific mRNA purification by translating ribosome affinity purification (TRAP). Nat Protoc 9, 1282–1291.

Heiman, M., Schaefer, A., Gong, S., Peterson, J.D., Day, M., Ramsey, K.E., Suarez-Farinas, M., Schwarz, C., Stephan, D.A., Surmeier, D.J., et al. (2008). A translational profiling approach for the molecular characterization of CNS cell types. Cell 135, 738–748.

Higashimori, H., Schin, C.S., Chiang, M.S., Morel, L., Shoneye, T.A., Nelson, D.L., and Yang, Y. (2016). Selective Deletion of Astroglial FMRP Dysregulates Glutamate Transporter GLT1 and Contributes to Fragile X Syndrome Phenotypes In Vivo. J Neurosci 36, 7079–7094.

Ikeuchi, K., and Inada, T. (2016). Ribosome-associated Asc1/RACK1 is required for endonucleolytic cleavage induced by stalled ribosome at the 3’ end of nonstop mRNA. Sci Rep 6, 28234.

Jannot, G., Bajan, S., Giguere, N.J., Bouasker, S., Banville, I.H., Piquet, S., Hutvagner, G., and Simard, M.J. (2011). The ribosomal protein RACK1 is required for microRNA function in both C. elegans and humans. EMBO Rep 12, 581–586.

Johnson, A.G., Lapointe, C.P., Wang, J., Corsepius, N.C., Choi, J., Fuchs, G., and Puglisi, J.D. (2019). RACK1 on and off the ribosome. RNA 25, 881–895.

Juszkiewicz, S., and Hegde, R.S. (2017). Initiation of Quality Control during Poly(A) Translation Requires Site-Specific Ribosome Ubiquitination. Mol Cell 65, 743–750 e744.

Juszkiewicz, S., Speldewinde, S.H., Wan, L., Svejstrup, J.Q., and Hegde, R.S. (2020). The ASC-1 Complex Disassembles Collided Ribosomes. Mol Cell 79, 603–614 e608.

Kershner, L., and Welshhans, K. (2017a). RACK1 is necessary for the formation of point contacts and regulates axon growth. Dev Neurobiol 77, 1038–1056.

Kershner, L., and Welshhans, K. (2017b). RACK1 regulates neural development. Neural Regen Res 12, 1036–1039.

Kharade, S.V., Kurata, H., Bender, A.M., Blobaum, A.L., Figueroa, E.E., Duran, A., Kramer, M., Days, E., Vinson, P., Flores, D., et al. (2018). Discovery, Characterization, and Effects on Renal Fluid and Electrolyte Excretion of the Kir4.1 Potassium Channel Pore Blocker, VU0134992. Mol Pharmacol 94, 926–937.

Kucheryavykh, Y.V., Kucheryavykh, L.Y., Nichols, C.G., Maldonado, H.M., Baksi, K., Reichenbach, A., Skatchkov, S.N., and Eaton, M.J. (2007). Downregulation of Kir4.1 inward rectifying potassium channel subunits by RNAi impairs potassium transfer and glutamate uptake by cultured cortical astrocytes. Glia 55, 274–281.

Kuroha, K., Akamatsu, M., Dimitrova, L., Ito, T., Kato, Y., Shirahige, K., and Inada, T. (2010). Receptor for activated C kinase 1 stimulates nascent polypeptide-dependent translation arrest. EMBO Rep 11, 956–961.

Lee, Y., Messing, A., Su, M., and Brenner, M. (2008). GFAP promoter elements required for region-specific and astrocyte-specific expression. Glia 56, 481–493.

Lin, Y.J., Huang, L.H., and Huang, C.T. (2013). Enhancement of heterologous gene expression in Flammulina velutipes using polycistronic vectors containing a viral 2A cleavage sequence. PLoS One 8, e59099.

MacVicar, B.A., Feighan, D., Brown, A., and Ransom, B. (2002). Intrinsic optical signals in the rat optic nerve: role for K(+) uptake via NKCC1 and swelling of astrocytes. Glia 37, 114–123.

Majzoub, K., Hafirassou, M.L., Meignin, C., Goto, A., Marzi, S., Fedorova, A., Verdier, Y., Vinh, J., Hoffmann, J.A., Martin, F., et al. (2014). RACK1 controls IRES-mediated translation of viruses. Cell 159, 1086–1095.

Mauro, V.P., and Matsuda, D. (2016). Translation regulation by ribosomes: Increased complexity and expanded scope. RNA Biol 13, 748–755.

Mazare, N., Oudart, M., Cheung, G., Boulay, A.C., and Cohen-Salmon, M. (2020a). Immunoprecipitation of Ribosome-Bound mRNAs from Astrocytic Perisynaptic Processes of the Mouse Hippocampus. STAR Protoc 1, 100198.

Mazare, N., Oudart, M., and Cohen-Salmon, M. (2021). Local translation in perisynaptic and perivascular astrocytic processes - a means to ensure astrocyte molecular and functional polarity? J Cell Sci 134.

Mazare, N., Oudart, M., Moulard, J., Cheung, G., Tortuyaux, R., Mailly, P., Mazaud, D., Bemelmans, A.P., Boulay, A.C., Blugeon, C., et al. (2020b). Local Translation in Perisynaptic Astrocytic Processes Is Specific and Changes after Fear Conditioning. Cell Rep 32, 108076.

McGough, N.N., He, D.Y., Logrip, M.L., Jeanblanc, J., Phamluong, K., Luong, K., Kharazia, V., Janak, P.H., and Ron, D. (2004). RACK1 and brain-derived neurotrophic factor: a homeostatic pathway that regulates alcohol addiction. J Neurosci 24, 10542–10552.

Miller, P.M., Folkmann, A.W., Maia, A.R., Efimova, N., Efimov, A., and Kaverina, I. (2009). Golgi-derived CLASP-dependent microtubules control Golgi organization and polarized trafficking in motile cells. Nat Cell Biol 11, 1069–1080.

Natchiar, S.K., Myasnikov, A.G., Kratzat, H., Hazemann, I., and Klaholz, B.P. (2017). Visualization of chemical modifications in the human 80S ribosome structure. Nature 551, 472–477.

Nielsen, M.H., Flygaard, R.K., and Jenner, L.B. (2017). Structural analysis of ribosomal RACK1 and its role in translational control. Cell Signal 35, 272–281.

Nilsson, J., Sengupta, J., Frank, J., and Nissen, P. (2004). Regulation of eukaryotic translation by the RACK1 protein: a platform for signalling molecules on the ribosome. EMBO Rep 5, 1137–1141.

Nwaobi, S.E., Cuddapah, V.A., Patterson, K.C., Randolph, A.C., and Olsen, M.L. (2016). The role of glial-specific Kir4.1 in normal and pathological states of the CNS. Acta Neuropathol 132, 1–21.

Olsen, M.L., and Sontheimer, H. (2008). Functional implications for Kir4.1 channels in glial biology: from K+ buffering to cell differentiation. J Neurochem 107, 589–601.

Oudart, M., Tortuyaux, R., Mailly, P., Mazare, N., Boulay, A.C., and Cohen-Salmon, M. (2020). AstroDot - a new method for studying the spatial distribution of mRNA in astrocytes. J Cell Sci 133.

Perez-Riverol, Y., Csordas, A., Bai, J., Bernal-Llinares, M., Hewapathirana, S., Kundu, D.J., Inuganti, A., Griss, J., Mayer, G., Eisenacher, M., et al. (2019). The PRIDE database and related tools and resources in 2019: improving support for quantification data. Nucleic Acids Res 47, D442–D450.

Pilaz, L.J., Lennox, A.L., Rouanet, J.P., and Silver, D.L. (2016). Dynamic mRNA Transport and Local Translation in Radial Glial Progenitors of the Developing Brain. Curr Biol 26, 3383–3392.

Poullet, P., Carpentier, S., and Barillot, E. (2007). myProMS, a web server for management and validation of mass spectrometry-based proteomic data. Proteomics 7, 2553–2556.

Radomska, K.J., Halvardson, J., Reinius, B., Lindholm Carlstrom, E., Emilsson, L., Feuk, L., and Jazin, E. (2013). RNA-binding protein QKI regulates Glial fibrillary acidic protein expression in human astrocytes. Hum Mol Genet 22, 1373–1382.

Ran, F.A., Hsu, P.D., Wright, J., Agarwala, V., Scott, D.A., and Zhang, F. (2013). Genome engineering using the CRISPR-Cas9 system. Nat Protoc 8, 2281–2308.

Risher, W.C., Andrew, R.D., and Kirov, S.A. (2009). Real-time passive volume responses of astrocytes to acute osmotic and ischemic stress in cortical slices and in vivo revealed by two-photon microscopy. Glia 57, 207–221.

Rollins, M.G., Jha, S., Bartom, E.T., and Walsh, D. (2019). RACK1 evolved species-specific multifunctionality in translational control through sequence plasticity within a loop domain. J Cell Sci 132.

Russo, A., Scardigli, R., La Regina, F., Murray, M.E., Romano, N., Dickson, D.W., Wolozin, B., Cattaneo, A., and Ceci, M. (2017). Increased cytoplasmic TDP-43 reduces global protein synthesis by interacting with RACK1 on polyribosomes. Hum Mol Genet 26, 1407–1418.

Sakers, K., Lake, A.M., Khazanchi, R., Ouwenga, R., Vasek, M.J., Dani, A., and Dougherty, J.D. (2017). Astrocytes locally translate transcripts in their peripheral processes. Proc Natl Acad Sci U S A 114, E3830–E3838.

Sakers, K., Liu, Y., Llaci, L., Lee, S.M., Vasek, M.J., Rieger, M.A., Brophy, S., Tycksen, E., Lewis, R., Maloney, S.E., et al. (2021). Loss of Quaking RNA binding protein disrupts the expression of genes associated with astrocyte maturation in mouse brain. Nat Commun 12, 1537.

Seifert, G., Huttmann, K., Binder, D.K., Hartmann, C., Wyczynski, A., Neusch, C., and Steinhauser, C. (2009). Analysis of astroglial K+ channel expression in the developing hippocampus reveals a predominant role of the Kir4.1 subunit. J Neurosci 29, 7474–7488.

Sibille, J., Dao Duc, K., Holcman, D., and Rouach, N. (2015). The neuroglial potassium cycle during neurotransmission: role of Kir4.1 channels. PLoS Comput Biol 11, e1004137.

Sibille, J., Pannasch, U., and Rouach, N. (2014). Astroglial potassium clearance contributes to short-term plasticity of synaptically evoked currents at the tripartite synapse. J Physiol 592, 87–102.

Sitron, C.S., Park, J.H., and Brandman, O. (2017). Asc1, Hel2, and Slh1 couple translation arrest to nascent chain degradation. RNA 23, 798–810.

Srinivasan, R., Lu, T.Y., Chai, H., Xu, J., Huang, B.S., Golshani, P., Coppola, G., and Khakh, B.S. (2016). New Transgenic Mouse Lines for Selectively Targeting Astrocytes and Studying Calcium Signals in Astrocyte Processes In Situ and In Vivo. Neuron 92, 1181–1195.

Sundaramoorthy, E., Leonard, M., Mak, R., Liao, J., Fulzele, A., and Bennett, E.J. (2017). ZNF598 and RACK1 Regulate Mammalian Ribosome-Associated Quality Control Function by Mediating Regulatory 40S Ribosomal Ubiquitylation. Mol Cell 65, 751–760 e754.

Takeuchi, A., Takahashi, Y., Iida, K., Hosokawa, M., Irie, K., Ito, M., Brown, J.B., Ohno, K., Nakashima, K., and Hagiwara, M. (2020). Identification of Qk as a Glial Precursor Cell Marker that Governs the Fate Specification of Neural Stem Cells to a Glial Cell Lineage. Stem Cell Reports.

The, M., MacCoss, M.J., Noble, W.S., and Kall, L. (2016). Fast and Accurate Protein False Discovery Rates on Large-Scale Proteomics Data Sets with Percolator 3.0. J Am Soc Mass Spectrom 27, 1719–1727.

Thompson, M.K., Rojas-Duran, M.F., Gangaramani, P., and Gilbert, W.V. (2016). The ribosomal protein Asc1/RACK1 is required for efficient translation of short mRNAs. Elife 5.

Valot, B., Langella, O., Nano, E., and Zivy, M. (2011). MassChroQ: a versatile tool for mass spectrometry quantification. Proteomics 11, 3572–3577.

Vedrenne, C., Klopfenstein, D.R., and Hauri, H.P. (2005). Phosphorylation controls CLIMP-63-mediated anchoring of the endoplasmic reticulum to microtubules. Mol Biol Cell 16, 1928–1937.

Wang, H., and Friedman, E. (2001). Increased association of brain protein kinase C with the receptor for activated C kinase-1 (RACK1) in bipolar affective disorder. Biol Psychiatry 50, 364–370.

Wenzel, H.J., Murray, K.D., Haify, S.N., Hunsaker, M.R., Schwartzer, J.J., Kim, K., La Spada, A.R., Sopher, B.L., Hagerman, P.J., Raske, C., et al. (2019). Astroglial-targeted expression of the fragile X CGG repeat premutation in mice yields RAN translation, motor deficits and possible evidence for cell-to-cell propagation of FXTAS pathology. Acta Neuropathol Commun 7, 27.

Xu, X., Yang, X., Xiong, Y., Gu, J., He, C., Hu, Y., Xiao, F., Chen, G., and Wang, X. (2015). Increased expression of receptor for activated C kinase 1 in temporal lobe epilepsy. J Neurochem 133, 134–143.

