## Supplementary material for "Translational regulation by RACK1 in astrocytes represses KIR4.1 expression and regulates neuronal activity": Key resource table

**KEY RESOURCES TABLE**

| REAGENT or RESOURCE | SOURCE | Dilution | IDENTIFIER |
|---|---|---|---|
| Antibodies | | | |
| Chicken polyclonal anti-GFP | Aves | 1:500e (WB) | Cat#GFP-1020<br>RRID:AB_10000240 |
| Rabbit polyclonal anti-GLT1 | Frontier Institute | 1:10,000e (WB) | Cat#Rb-Af670<br>RRID:AB_2571718 |
| Rabbit polyclonal anti-KIR4.1 | Alomone labs | 1:500e (WB) | Cat#APC-035<br>RRID:AB_2040120 |
| Mouse monoclonal anti-RACK1 (B-3) | Santa Cruz Biotechnology | 1:500e (WB) | Cat#sc-17754<br>RRID:AB_2247471 |
| Mouse monoclonal anti-RACK1 | BD transduction laboratories | 1:500e (IF) | Cat#610177<br>RRID:AB_397576 |
| Rabbit monoclonal anti-RPS6 | Cell signaling Technology | 1:500e (WB) | Cat#2217<br>RRID:AB_331355 |
| Rabbit polyclonal anti-GFAP | Sigma-Aldrich | 1:500e (IF) | Cat#G9269<br>RRID:AB_477035 |
| Peroxidase AffiniPure Goat anti-chicken IgY (IgG) (H+L) | Jackson ImmunoResearch | 1:10,000e (WB) | Cat#103-035-155<br>RRID:AB_2337381 |
| Goat anti-rabbit IgG (H+L) – HRP | CliniSciences | 1:2,500e (WB) | Cat#CSA2115 |
| Goat anti-mouse IgG (H+L) – HRP | CliniSciences | 1:2,500e (WB) | Cat#CSA2108 |
| EasyBlot anti Mouse IgG (HRP) | GeneTex | 1:1,000e (WB) | Cat#GTX221667-01<br>RRID:AB10728926 |
| EasyBlot anti Rabbit IgG (HRP) | GeneTex | 1:1,000 (WB) | Cat#GTX221666-01<br>RRID:AB_10620421 |
| Goat anti-Rabbit IgG (H+L) Alexa Fluor 488 | Invitrogen | 1:1,000e (IF) | Cat#A-11034<br>RRID:AB_2576217 |
| Goat anti-Mouse IgG (H+L) Alexa Fluor 555 | Invitrogen | 1:1,000e (IF) | Cat#A-21424<br>RRID:AB_141780 |
| Mouse IgG Isotype Control | Thermo Fisher Scientific | 25 ug | Cat#10400C<br>RRID:AB_2532980 |
| FISH Probes | | | |
| RNAscope Probe Mm-Gnb2l1-E1-E3 | Bio-Techne | Cat#443621 | NM_008143.3 |
| qPCR probes | | | |
| Slc1a2 | Thermofisher | Mm01275814_m1 | NM_001077514.3 |
| Gjb6 | Thermofisher | Mm00433661_s1 | NM_001010937.2 |
| Gja1 | Thermofisher | Mm01179639_s1 | NM_010288.3 |
| Slc1a3 | Thermofisher | Mm00600697_m1 | NM_148938.3 |
| Aqp4 | Thermofisher | Mm00802131_m1 | NM_009700.2 |
| Kcnj10 | Thermofisher | Mm00445028_m1 | NM_001039484.1 |
| Gnb2l1 | Thermofisher | Mm01291968_g1 | NM_008143.3 |
| rRNA18S | Thermofisher | Mm03928990_g1 | NR_003278.3 |
| Primers | | | |
| **Mouse genotyping** | | | |
| Cre recombinase transgene forward | CTTCAACAGGTGCCTTCCA | | |
| Cre recombinase transgene reverse | GGCAAACGGACAGAAGCA | | |
| Internal positive control forward | CTGTCCCTGTATGCCTCTGG | | |
| Internal positive control reverse | AGATGGAGAAAGGACTAGGCTACA | | |

| | |
|---|---|
| Floxed Rack1 forward | GAACGCTGCGCCTCTGGGATCTCACAA |
| Floxed Rack1 reverse | AGTCCACACGTGGCTTAGAACCTA |
| **Brain Rack1 deletion** | |
| Forward | GACCTGACACGCACTTCTGA |
| Reverse | CAGTCCACACGTGGCTTAGA |
| ***Kcnj10* 5'UTR PCR fragments** | |
| 1-146 For | AAGCTAGCACAAAGTTTGGCTTCGGCACTGCAGAGGG |
| 1-146 Rev | AAGCTAGCACAGGCAGGGCCAAGACTCT |
| 127-242 For | AAGCTAGCAGAGTCTTGGCCCTGCCTGT |
| 127-242 Rev | AAGCTAGCCTTGGCTCGAAGGTGAAGTGGCTGCTCTG |
| 95-242 For | AAGCTAGCCAAACCCTTATCTGATTCCAG |
| 95-242 Rev | AAGCTAGCCTTGGCTCGAAGGTGAAGTGGCTGCTCTG |
| 95-191 For | AAGCTAGCCAAACCCTTATCTGATTCCAG |
| 95-191 Rev | AAGCTAGCAGATGAACAGACGGGGCGG |
| 1-191 For | AAGCTAGCACAAAGTTTGGCTTCGGCACTGCAGAGGG |
| 1-191 Rev | AAGCTAGCAGATGAACAGACGGGGCGG |
| 181-242 For | AAGCTAGCTCTGTTCATCTGTCCCGCT |
| 181-242 Rev | AAGCTAGCCTTGGCTCGAAGGTGAAGTGGCTGCTCTG |
| **CRISPR/cas9-mediated genome editing** | |
| human *Rack1*/*Gnb2l1 For* | CACCGTGTCAACCGCACGTCTATGC |
| human *Rack1*/*Gnb2l1 Rev* | AAACGCATAGACGTGCGGTTGACAC |
| **Cloning sequences** | |
| Linker BspEI1-P2A-Flag-EcoRI-NotI-KpnI | AAGTCCGGAGGCAGCGGCGCCACCAACTTTTCCCTGCT<br>CAAGCAGGCCGGCGACGTGGAAGAGAATCCCGGCCCC<br>GGTACTGACTACAAAGACCATGACGGTGATTATAAAG<br>ATCATGACATCGATTACAAGGATGACGATGACAAGGA<br>ATTCGCGGCCGCAGGTACCGGA |
| mouse *Kcnj10* CDS gBlock - EcoRI/NotI | AAAGAATTCACGTCGGTCGCTAAGGTCTATTACAGTCA<br>GACGACTCAGACAGAGAGCCGCCCCCTAGTGGCCCCA<br>GGAATACGCCGGAGGAGGGTCCTCACGAAAGACGGCC<br>GGAGCAATGTGAGAATGGAGCACATTGCTGACAAACG<br>TTTCCTCTACCTCAAGGATCTATGGACGACCTTCATTG<br>ACATGCAATGGCGCTACAAGCTTCTGCTCTTCTCTGCA<br>ACCTTTGCAGGCACGTGGTTCCTCTTTGGTGTGGTGTG<br>GTATCTGGTAGCTGTGGCCCATGGGGACCTGTTGGAGC<br>TGGGACCTCCTGCCAACCACACGCCTTGTGTGGTGCAG<br>GTGCACACGCTCACCGGAGCCTTCCTCTTCTCCCTGGA<br>ATCCCAGACCACCATCGGCTATGGCTTCCGCTACATCA<br>GTGAGGAATGCCCACTGGCCATCGTGCTTCTTATTGCG<br>CAGCTGGTGCTCACCACCATTCTGGAAATCTTCATCAC<br>AGGTACCTTCCTTGCAAAGATTGCCCGGCCTAAGAAGA<br>GGGCCGAGACGATCCGCTTCAGCCAGCATGCCGTTGTG<br>GCTTCCCATAACGGGAAGCCTTGCCTTATGATCCGGGT<br>TGCCAATATGCGGAAGAGTCTCCTCATTGGATGCCAGG<br>TGACAGGCAAACTGCTTCAAACGCACCAGACAAAGGA<br>GGGTGAGAATATTCGGCTCAACCAGGTCAACGTGACTT<br>TCCAAGTAGACACAGCCTCAGACAGCCCCTTTCTCATC<br>CTACCCCTGACTTTCTACCACGTGGTAGATGAGACCAG<br>CCCCTTAAAAGATCTCCCGCTCCGCAGTGGGGAGGGG |

| | |
|---|---|
| | GACTTTGAGCTGGTGCTGATCCTGAGTGGGACAGTGGA<br>GTCCACCAGTGCCACCTGCCAAGTTCGCACTTCCTACC<br>TACCGGAGGAGATCCTCTGGGGTTACGAGTTCACGCCT<br>GCGATCTCACTGTCAGCCAGTGGCAAATACATAGCTGA<br>CTTCAGCCTTTTCGACCAGGTTGTGAAAGTGGCATCTC<br>CCAGTGGTCTCCGCGATAGCACCGTACGCTATGGAGAC<br>CCCGAGAAGCTCAAGTTGGAGGAGTCATTAAGAGAGC<br>AAGCTGAAAAGGAAGGCAGTGCCCTTAGTGTGCGCAT<br>CAGCAACGTCGCGGCCGCAAA |
| mouse *Kcnj10* 5'UTR#1 (NM_001039484.1) – NheI/NheI | AAGCTAGCACAAAGTTTGGCTTCGGCACTGCAGAGGG<br>AGGGGGCGGCCCGGCCCGGCCCAACTCTGCCCCCGGC<br>CGGCCCGACCCCGGCCCCGGCCCCCGGACAAACCCTTA<br>TCTGATTCCAGCTCCGGGTTTAAGAGTCTTGGCCCTGC<br>CTGTCGCACAGCTCCGCTCGCCGCTCCTGCCCGCCCCG<br>TCTGTTCATCTGTCCCGCTGCCCCGCGATTTATCAGAGC<br>AGCCACTTCACCTTCGAGCCAAGGCTAGCAA |
| mouse *Kcnj10* 5'UTR#2 (AB039879.1) – NheI/NheI | AAAGCTAGCCCGCGCACAGCTCCGCTCGCCGCTCCTGC<br>CCGCCCCGTCTGTTCATCTGTCCCGCTGCCCCGCGATTT<br>ATCAGAGCAGCCACTTCACCTTCGAGCCAAGGCTAGCA<br>AA |
| mouse *Kcnj10* 3'UTR (AB039879.1) – SalI/NotI | AAAGTCGACTATCCTGTTCTCTCCTCCCCAGCCCTCTGG<br>CCTTTTCCTCTTCCAGTGCTCTGGGAAAAGGATACAAT<br>CTGGGTTCGCTGGAGAGACCCTGAAGCACCCACCCTCA<br>ACTTCCCCTTAGCCCGGTGGCCTGTGAGCAGTCGGGGC<br>CTGCTGGAGCCCTTCCTTTTCCTCTCACTGTCCCCTTGT<br>ACCTTCCTCTACCACTACACCAATGTATATGACTCTTA<br>AGCCAGCTTGGGGGAAAGAGAGGGAAGATGAGGCTGA<br>CATGGCTTGGAAGGCCGCAGCCATGCTTGGAGATTCAC<br>ATTCAGAGGACCATGTGACTGGATGGATAGACTCCCCC<br>CCAAACGCCCACCACTAGAAAATTTGATGGCTTGAGAC<br>TGGAGCTTCCTCCTTTTCTCTTCCATCTAGCTAAGTTCC<br>CTGAGGCAGAACTCTCTCAGAGAGTCACTCTATGGGCT<br>CTGCCCTCAGAAGTACTGGGATCTGAAATCAGTTTTCC<br>CTGGGCTATCAAATCCTGAAGAAAGAAGCCAGAGTTT<br>GTCATGGTGTTTTAATTTCAGCTGGTAACTCCTGACAG<br>AGCCTGGACTCTTGGTCACCACCTAATTTTCCCTCTCCA<br>GTTCTCAAGTGGATCTTCTCAGCAAAGCCGGTTCTCTC<br>TCAGCTTCTGTTTCTGTAGAACTGGGGTTGACTCGAGA<br>AACACAAGCAGCCACCTTTCCTGCACCCATCCTGGATT<br>CTGAGAAATTCAATTTTGGAAATACTGTCTAATACTCT<br>GGCCCTGTCCTTTGGGCAGCTCTGTCCTGAAAAGATAT<br>GTTTTAGTGTTTCCTGGGAATAGGAAGACCTTAACTCG<br>TGCCGCGGCCGCAAA |

| Experimental Models: Organisms/Strains | | |
|---|---|---|
| Mouse: C57BL/6J wildtype | Janvier labs | SC-C57J-M |
| Mouse: Tg(Aldh1l1-eGFP/Rpl10a) JD130Htz BacTRAP | Nathaniel Heintz's laboratory (Rockefeller University, New York City, NY) | MGI:5496674 www.bactrap.org |

| | | | |
|---|---|---|---|
| Mouse: B6J.Cg-Rack1$^{tm1.1Cart}$/Mmucd Rack1$^{fl/fl}$ | MMRRC | MGI: 6343619 | |
| Mouse: B6N.FVB-Tg(Aldh1l1-cre/ERT2)1Khakh/J Aldh1l1-CreERT2 | The Jackson Laboratory | MGI: 5806593<br>JAX: 029655 | |
| Software and Algorithms | | | |
| ImageJ | Schindelin et al., Nature Methods, 2012 | ImageJ 1.53f51 | https://imagej.nih.gov/ij/ |
| AstroDot plugin for ImageJ | Our laboratory | | Oudart et al., 2020 |
| GraphPad Prism | GraphPad Software, Inc. | Version 8.0.3 (263) | |
| Virus Strains | | | |
| Serotype 9 AAV GfaABC$_1$D:TdTOMATO | | | |
| Deposited Data | | | |
| Mass Spectrometry data on ProteomeXchange via PRIDE database | PXD033121 | | HXvtnu9v |
| Chemicals, Peptides, and Recombinant Proteins | | | |
| VU 0134992 KIR4.1 Blocker | Biotechne | 30 uM | Cat#6877 |
